# Engineering the Self-assembly of Bacterial Microcompartment Shell Proteins via Charged Mutations

**DOI:** 10.64898/2026.01.29.702620

**Authors:** Annie Gomez, Behzad Mehrafrooz, Curt Waltmann, Carolyn E. Mills, Nolan W. Kennedy, Jacob B. Miller, Danielle Tullman-Ercek, Monica Olvera de la Cruz

**Affiliations:** Department of Materials Science and Engineering, Northwestern University, Evanston, IL 60208; Center for Computation & Theory of Soft Materials, McCormick School of Engineering, Northwestern University, Evanston, IL 60208; Center for Synthetic Biology, Northwestern University, Evanston, IL, 60208, USA; Department of Chemical and Biological Engineering, Northwestern University, Evanston, IL, 60208, USA; Interdisciplinary Biological Sciences, Northwestern University, Evanston, IL, 60208, USA; Department of Chemistry, Northwestern University, Evanston, IL, 60208, USA

## Abstract

Protein self-assembly is a fundamental biological process of great importance for the design and synthesis of biomaterials. Developing the ability to precisely manipulate protein assembly would greatly expand both our understanding of the process and our biotechnological capabilities. Within bacteria, proteins that self-organize to form bacterial microcompartments (MCPs) offer an excellent model system for studying protein self-assembly and advancing biomaterial design capabilities. MCPs consist of irregular polyhedral shells that encase an enzyme core, functioning as enzymatic nanoreactors. In isolation, the abundant shell proteins of the 1,2-propanediol utilization (Pdu) MCP, PduA and PduJ, have a high propensity to self-assemble into tubular structures, analogous in form to carbon nanotubes. Here, we design and characterize tubular structures formed by hexameric PduA and PduJ proteins. We demonstrate that altering hexamer charge offers a systematic strategy for modulating the higherorder assembly of PduA and PduJ across multiple contexts by integrating molecular dynamics simulations with heterologous overexpression, cell-free, and *in vivo Salmonella enterica* serovar Typhimurium LT2 experiments. First, using molecular simulation, we find that tube chirality and radius play critical roles in determining structural stability and flexibility. Next, overexpression and cell-free experiments show that increasing the overall negative charge of assembling subunits consistently promotes self-assembly into tubular structures. We find that this holds true in the native MCP system, as these same mutations promote the formation of tubular MCP structures in *S. enterica* LT2. Our results collectively reveal that both electrostatic interactions and fields generated by charges on proteins can be leveraged to control protein-based nanostructures.

## Introduction

Elucidating the key fundamental principles of biomolecular self-assembly is essential to understanding biological systems and designing biomimetic materials.^1–3^ Protein assemblies are a critical subset of such biomolecular assemblies in which sequence-defined protein building blocks autonomously organize into nanostructures. ^4,5^ The resultant nanostructures facilitate functions such as catalysis via compartmentalization, as is the case for bacterial organelles called bacterial microcompartments (MCPs),^6,7^ which support enzymatic transformations that promote bacterial survival, often in harsh or nutrient-poor environments, by encapsulating specific enzymes in a protein shell.^8–10^ MCP shells are composed entirely of hexameric, trimeric, and pentameric proteins that assemble into regular or irregular polyhedra (75−200 nm in diameter)^11–14^ where the appearance of irregular polyhedral MCPs can be explained through modeling of multicomponent crystalline shell assembly.^5,15,16^ Interestingly, while MCPs can appear geometrically similar to viral capsids,^17^ MCP shells encapsulate specific metabolic enzymes^12^ rather than genetic material.^18^

The well-characterized 1,2-propanediol utilization (Pdu) MCP in *Salmonella enterica* serovar Typhimurium LT2^19^ has served as a model system for interrogating the assembly and biological functions of metabolic MCPs, which include toxic intermediate sequestration,^8^ and enzyme colocalization to control cofactor recycling and pathway flux.^20^ The most abundant proteins in the Pdu MCP shell are PduA and PduJ, which are hexameric bacterial microcompartment domain (BMC-H) proteins.^21,22^ Each BMC-H hexamer consists of six identical protein monomers that assemble into flat or bowl-shaped six-sided structures, which then tile together to form the MCP shell’s facets in a honeycomb-like pattern that is reminiscent of graphene nanosheets.^23,24^ Interestingly, upon overexpression of PduA or PduJ in a non-native host, *Escherichia coli*, both proteins self-organize into nanotube structures with geometries reminiscent of carbon nanotubes, where tube lengths range from hundreds of nanometers to hundreds of micrometers. ^21,25^ These nanotubes hold great potential for future applications, including molecular encapsulation, targeted therapeutic delivery, and catalysis.^23,26,27^

Although PduJ and PduA are often considered functionally redundant in shell assembly and share ∼80% primary sequence identity,^21,28^ their divergent behaviors outside the native MCP environment highlight distinct self-assembly properties.^29,30^ These differences underscore that even subtle variations in assembling subunits can translate into variable assembly outcomes, driven in part by distinct intermolecular interactions. A key difference between PduA and PduJ is the overall net charge of their hexameric units—the PduJ hexamer has an overall net charge of −6 e, while the PduA hexamer is overall charge neutral. Previous studies indicate that charge is implicated in MCP assembly, ^5,31^ and that altering residues involved in hydrogen-bonding interactions at interfacial junctions can modulate overall MCP geometry.^5^ Here, we hypothesize that single point mutations that alter hexamer charge influence assembly, even when these charges are far away from the interfacial junction points. This is reasonable because proteins have a much lower dielectric constant (*E^p^* ≈ 3)^32^ than bulk water (*E*^bulk^ ≈ 80), which strengthens the electrostatic interactions (*U*) near the protein surface (*U* ∝ 1*/E^w^* with *E^w^* ≈ 7).^33^ Moreover, electric fields can propagate largely unscreened through the protein, which allows distal charge mutations to influence interactions over distances up to 10 nm, as demonstrated via single charge mutations in the SARS-COV-2 spike protein.^34^

In this work, we investigate the effects of three charge-altering point mutations at residues that differ between PduA and PduJ that reverse polarity, from positive to negative and *vice versa*. We examined the impact of these mutations on the self-assembly behavior of PduA and PduJ across *in vivo* and *in vitro* systems, and used molecular simulations to understand the intermolecular interactions that determine whether charge mutations in PduA and PduJ lead to symmetric morphological changes. These PduA and PduJ variants are investigated in several different contexts in this study, including the co-assembly with other native MCP shell proteins in *S. enterica*, the self-assembly upon overexpression in *E. coli*, and the self-assembly upon cell-free protein expression. Heterologous overexpression in *E. coli* and cell-free protein expression experiments reveal that surface charge residues located away from the hexameric junctions in PduA and PduJ significantly influence their assembly, promoting the formation of nanotube structures. Incorporating these charge variants into the *pdu* operon in *S. enterica* results in a higher instance of tubular MCP structures compared to the wild type (WT), but in less abundance than *in vitro*. Atomistic simulations are used to identify the conformations and intermolecular interactions responsible for PduA and PduJ nanotube assembly. Together, our findings show that charge-based variants can be leveraged to design different assembled geometries *in vitro* and *in vivo*, providing essential guidance for engineering these protein assemblies across a wide range of applications, from pathway encapsulation, to enzyme scaffolding.^35,36^

## Results and Discussion

PduA and PduJ exhibit high sequence similarity (82.9%) and high structural similarity (template modeling (TM) score of 0.96/1.0). ^37,38^ Interestingly, the residues that differ between PduA and PduJ include several charged amino acids, leading to a significant difference in net charge of the assembled hexamer. Specifically, PduA carries charged residues at positions E_4_, R_66_, and K_86_, while the corresponding positions in PduJ (shifted by one residue due to differences at the N terminus) are occupied by neutral residues N_3_, S_65_, and A_85_, respectively (Figure 1a). We hypothesize that this charge difference influences their assembly by modulating electrostatic interactions. These charge differences are all positioned at different locations in the hexamer. As shown in Figure 1b, residues N_3_ in PduJ and E_4_ in PduA are positioned near the center of the concave face. In contrast, while A_85_ in PduJ and K_86_ in PduA are also on the concave face of the hexamer, these residues are much closer to the hexamer-hexamer interface.^39^ Finally, S_65_ in PduJ and R_66_ in PduA are located near the center, again, away from the interface, on the convex face of the hexamer.

**Figure 1:**
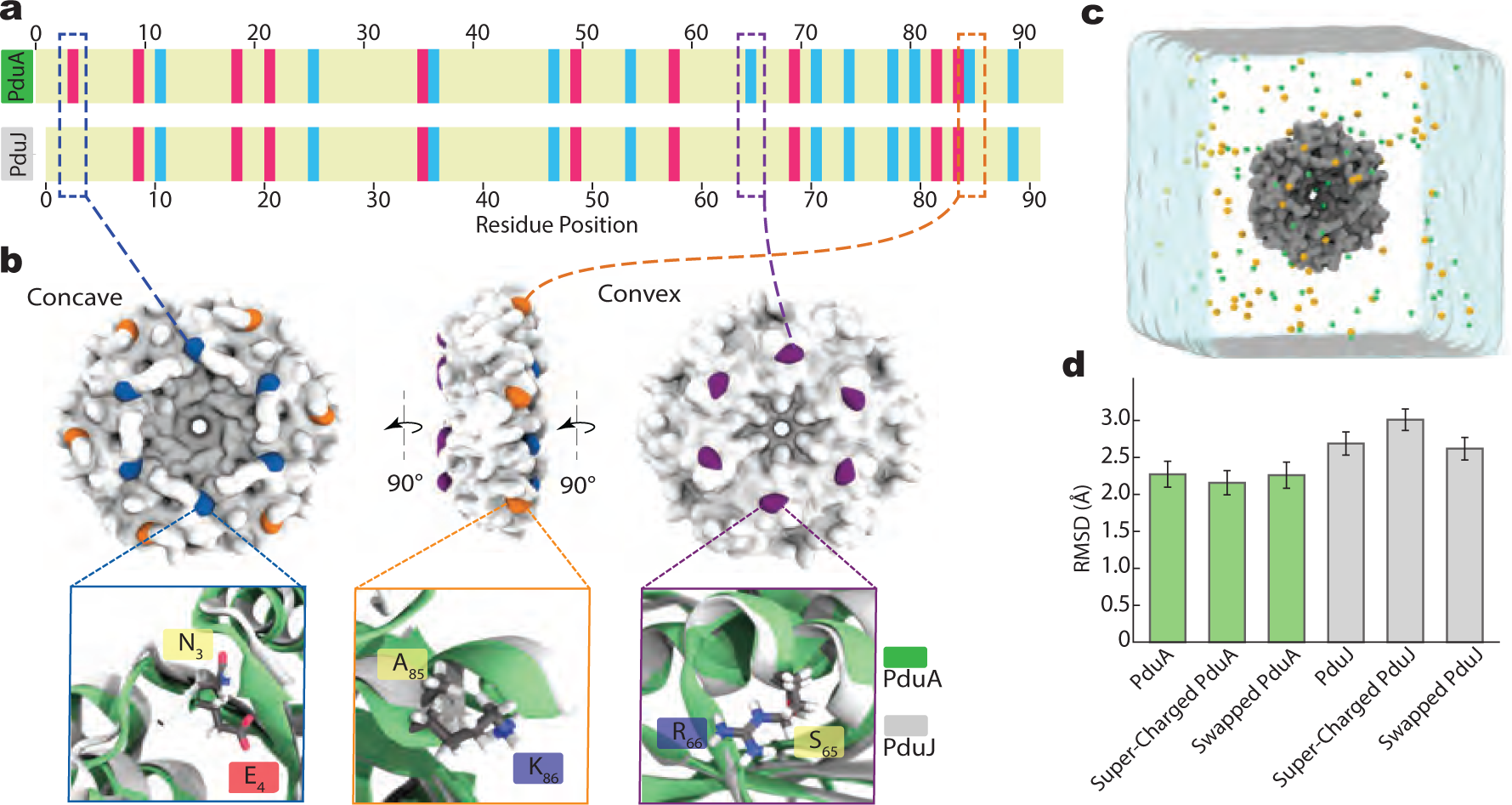
PduA and PduJ charged residues profiles. **a,** Residue-based alignment of PduA and PduJ amino acid sequence. The two-residue length difference results in a numbering offset, which causes PduJ to appear shifted relative to PduA in the alignment. The charge differences at positions 4, 66, and 86 in PduA (corresponding to 3, 65, and 85 in PduJ) are highlighted with dashed borders. Positively charged residues are shown in blue, negatively charged in pink, and neutral residues in yellow. **b,** Surface representations of concave, side, and convex view of PduA/J hemaxer (top row) and overlaid cartoon representation of the hexamers PduA (green) and PduJ (gray) at the differently charged residues (bottom row). Negatively, positively, neutral residues are labeled in red, blue and yellow, respectively. Sidechains are shown as sticks. **c,** All-atom model of a PduA/J hexamer (gray surface) submerged in a 14 14 14 nm^3^ water box (semi-transparent surface) containing 100 mM solution of NaCl (yellow and green spheres). **d,** RMSD of heavy atoms in protein’s secondary structure compared to the crystal structure coordinates in unrestrained simulations for PduA/J wildtype and mutant variants. Data show averages over ~370 ns trajectory and error bars are the standard error of the mean from four 90 ns sampling windows.

To investigate how these charge differences affect PduA and PduJ assembly, we studied two variants of each protein. The first variant of each protein swaps the amino acids at these residues in PduA and PduJ to the amino acids of PduJ and PduA, respectively, directly exchanging the PduA and PduJ charge profiles. These variants are herein called “swapped” variants. Specifically, swapped PduA refers to PduA−E_4_N−R_66_S−K_86_A, and swapped PduJ refers to PduJ−N_3_E−S_65_R−A_85_K. Thus, the “swapped PduA” hexameric unit has a −6 e net charge (compared to the net neutral charge of the wild type PduA hexamer), while the “swapped PduJ” hexameric unit has a net neutral charge (compared to the −6 e net charge of the wild type PduJ hexamer). The second variant of each protein, referred to as the “supercharged” variant, features mutations of the residues of interest to charged amino acids with charge opposite to that of the corresponding residues in PduA. Specifically, the first of these residues, which is a glutamic acid (E) in wild type PduA was mutated to a positively charged lysine (K), and the second and third of these residues, which are positively charged arginine (R) and lysine (K) in wild type PduA, were both mutated to negatively charged glutamic acid (E). Both supercharged hexamer variants carry a net charge of −12 e. The specific protein modifications and corresponding charge annotations for the wild-type, swapped, and supercharged mutants are summarized in Table 1, for the specific genetic backgrounds used in different experiments, see SI table 1.

**Table 1:** Variants of PduA and PduJ and their charge properties.

| **Variant name** | **Charge-based mutation** | **Individual charge mutation $\Delta$** | **Hexamer net charge (e)** |
| --- | --- | --- | --- |
| **PduA variants** |  |  |  |
| WT PduA | N/A | N/A | 0 |
| Swapped PduA | E<sub>4</sub>N–R<sub>66</sub>S–K<sub>86</sub>A | +1, –1, –1 | –6 |
| Supercharged PduA | E<sub>4</sub>K–R<sub>66</sub>E–K<sub>86</sub>E | +2, –2, –2 | –12 |
| **PduJ variants** |  |  |  |
| WT PduJ | N/A | N/A | –6 |
| Swapped PduJ | N<sub>3</sub>E–S<sub>65</sub>R–A<sub>85</sub>K | –1, +1, +1 | 0 |
| Supercharged PduJ | N<sub>3</sub>K–S<sub>65</sub>E–A<sub>85</sub>E | +1, –1, –1 | –12 |

We assess how these mutations influence the structure of individual PduA and PduJ hexamers by performing all-atom molecular dynamics (MD) simulations on both the WT and variant hexamers. Such detailed structural changes are difficult to resolve experimentally, making MD simulations essential. Our simulation setup included a variant of each protein solvated in a 14 × 14 × 14 nm^3^ box of 100 mM NaCl in water (Figure 1c). The stability of the hexamers was quantified by computing the root mean square deviation (RMSD) of non-hydrogen atoms comprising the *β*-sheets and *α*-helices of the structure over the course of the 370 ns free equilibration simulation. The RMSD of the proteins’ secondary structures in all PduA simulations remained, on average, at ∼2.5 ^°^A (Figure 1d), while PduJ variants showed slightly higher RMSD (±0.5 ^°^A). This suggests that charge-based mutations do not disrupt the intramolecular interactions that stabilize the hexameric structure, and thus do not affect the overall structural integrity of the hexamers.

### Distinct Self-Assembly Behaviors of PduA and PduJ when Over-expressed in Non-Native Systems

We initially explored the impact of charge mutations on the self-assembly behavior of PduA and PduJ by overexpressing our variants in *E. coli*, a host that enables production of individual PduA or PduJ proteins independent of other MCP components.^21,23^ Cells expressing PduA and PduJ variants can self-assemble in cells, appearing primarily as long chains of adjoined cells (Figure 2a), which are physically linked due to self-assembled PduA or PduJ tubes within the cytoplasm that prevent cell division.^21^ These tubes are composed of hexameric PduA or PduJ subunits arranged side-by-side, indicating that the hexamers in the assembly exhibit curvature relative to each other to form the observed tubular morphology (Figure 2a—bottom row).

**Figure 2:**
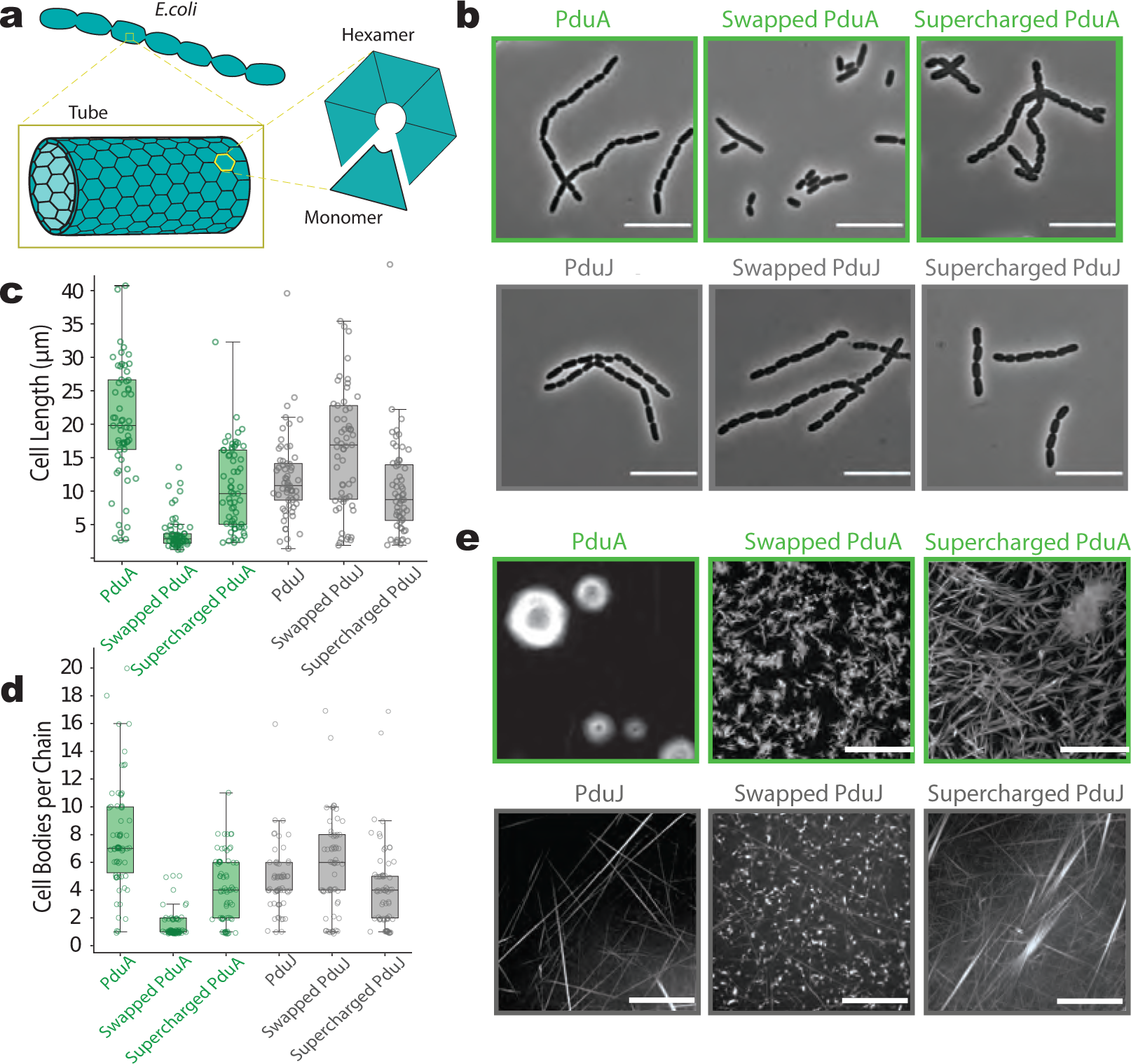
PduA and PduJ self-assemble into tubes differently*in vivo* and cell-free. **a,** Inset schematically shows *E. coli* cell containing PduA/J tubes. PduA/J tubes structures which run along the long axis of the cell body. These tubes are formed by rolling hexameric sheets with each hexamer consisting of six monomers. **b,** *E. coli* cells show WT and PduA/J mutants self-assembled into tubes when overexpressed (scale bar = 10 *µ*m). Images are captured using phase contrast microscopy. Hereafter, PduA variants are colored in green while PduJ in gray. **c,** Length of cells (defined as end-to-end distance per cell) induced for overexpression of PduA/J proteins. Box plots display distribution of cell lengths into quartiles and standard deviation across cell population. Measurements were obtained from three biological replicates. **d,** Fraction of linked or elongated cells in total population of cells for PduA/J and mutants overexpression. Cells were identified as cell bodies or “chains” if a clear cleavage furrow was present. An elongated cell, which is ¿ 10 *µ*m long cells with no cleavage furrow, is counted as a single cell. Counts were taken from three biological replicates(*>*80 cells counted per strain per replicate). **e,** Cell-free confocal microscopy imaging of PduA/J and mutants proteins show different structures are formed by different variants (scale bar = 10 *µ*m).

Upon overexpression of charge variants, we observed a range of cell division defects (Figure 2b), although all variants conferred a linked cell phenotype, suggesting that they were all assembly competent to some extent. Their degree of self-assembly is quantified by the cell length (defined as the end-to-end length of the cell chain, shown in Figure 2c) for each variant. Consistent with previous reports,^21^ cells expressing WT PduA were slightly longer than those expressing WT PduJ. Cells expressing WT PduA exhibited an average length of approximately 21.6 ± 12.6 *µ*m and typically formed chains of about eight cell bodies, while cells expressing WT PduJ averaged 11.9 ± 6.1*µ*m in length, forming shorter chains of about five cell bodies (Figure 2d). As a negative control, we also expressed assembly-deficient variants of PduA (K_26_A) and PduJ (K_25_A), which cannot form tubes,^21,39^ in contrast to assembly-competent WT proteins (see SI Figure 1). The expression of both WT proteins led to longer cells than their respective assembly-deficient mutants (on average, 2.68 ± 0.7 *µ*m for PduA−K_26_A and 2.60 ± 0.8 *µ*m for PduJ−K_25_A), as expected. These results confirm prior observations that while PduA and PduJ are structurally redundant in their native context, they do not behave identically when overexpressed in a heterologous host.^21,28^ The average cell length for swapped PduA decreased significantly to 3.65 ± 2.5 *µ*m compared to WT PduA and cells appeared primarily as single cell bodies, similar to strains expressing assembly-deficient variants of PduA and PduJ and normal *E. coli* morphology.^23^ This suggests that strains expressing swapped PduA do not behave like those expressing WT PduJ, despite the fact that swapped PduA and WT PduJ have the same net charge. In contrast, overexpression of swapped PduJ increases cell length relative to WT PduJ. Cells expressing swapped PduJ averaged 17.5 ± 11.2 *µ*m and formed chains of about six cell bodies, indicating enhanced self-assembly and a more severe division defect. These results suggest that the surface charge modulates self-assembly in a protein-specific manner.

To explore the impact of incorporating additional negative charge into the hexamers, we also expressed supercharged PduA and supercharged PduJ and assessed the resulting cell lengths. Supercharged PduA-expressing cells showed an intermediate phenotype compared to WT PduA and swapped PduA, averaging 10.6 ± 6.2 *µ*m in length and forming chains of approximately four cell bodies. This indicates self-assembly sufficient to interfere with cell division. Supercharged PduJ variants showed similar behavior, conferring an average cell length of 10.1 ± 6.9 *µ*m and chains of about four cell bodies, comparable to those seen in cells expressing WT PduJ. The observed and quantified differences in the average cell length among cells expressing PduA and PduJ variants provide strong evidence that charge-based mutations modulate the self-assembly behavior and consequently the severity of cell division defects in cells expressing these proteins (SI Figure 2).

To study how electrostatic charge influences the assembled morphologies of PduA and PduJ and their swapped and supercharged mutations in a simplified context, we carried out both experiments in cell-free systems and all-atom simulations of interactions between the proteins. Experimentally, cell-free protein synthesis (CFPS) enables direct observation of assemblies formed by proteins produced *in vitro*. This approach enables real-time detection by combining transcription–translation machinery, DNA, and fluorescent anti-FLAG antibodies in a single reaction, eliminating the need for purification.^29^ After confirming prior reports that WT PduA and WT PduJ self-assemble into hexagonal sheet-like structures and fibers (Figure 2e and SI Figure 3), ^29^ we proceeded to test our charge variants. Swapped PduJ assembles into long thin fibers that resemble the fibers observed in WT PduA; these fibers, however, co-exist with aggregates with a rice-like appearance and not with sheets. Supercharged PduJ assembles into extended fiber bundles when expressed in CFPS, supporting the idea that enhanced electrostatic interactions stabilize and promote axial growth and bundling, which is characteristic of self-assembled highly charged fibers.^40^ These observations suggest that supercharging the hexameric proteins enhances both elongation and directional bundling of assemblies formed upon expression in cell-free systems. We find that swapped PduA primarily forms short, bundled, anisotropic fiber-like structures. In contrast, similar to supercharged PduJ, supercharged PduA appears to form longer anisotropic fibril bundles. These fiber bundles can also form larger sheet-like structures, which can be seen in the upper right corner of the image (Figure 2e—2nd from the top right corner image). Both swapped and supercharged PduA variants form more fiber-like assemblies compared to WT PduA, which predominantly forms sheets. To probe the molecular driving force for this difference, we used all-atom molecular dynamics simulations to quantify the bending angle between pairs of WT PduJ, WT PduA, and supercharged PduA hexamers, computing the potential of mean force (PMF) as a function of inter-hexamer bending (see SI Figure 4). The PMF for WT PduA shows no strongly preferred bending angle, consistent with its tendency to form planar sheet-like assemblies.^41^ In contrast, supercharged PduA exhibits a narrower PMF minimum at non-zero bending angles, indicating a preference for curved, fiber-like assemblies. Together, these results demonstrate that electrostatic charge plays a critical role in modulating the curvature of PduA assemblies.

### In silico structure and dynamics of PduJ nanotubes

To gain a comprehensive molecular understanding of how the short range and electrostatic interactions drive tube assembly, we focus on WT PduJ, which, as noted above, is charged and exclusively forms tubes in cell free systems. To this end, we performed explicit-solvent, all-atom MD simulations of PduJ tubes with a range of diameters and chiralities. We borrow nomenclature from the carbon nanotube literature to describe the arrangement of hexamers within the tube, where the chirality, or helical twist of the hexameric sheet when it is rolled into a tube, can be either armchair, zigzag, or chiral. To build a molecular model of a PduJ nanotube, then, the tube diameter and chirality must be specified. While TEM analyses report an average diameter of ∼ 20 nm for PduJ tubes,^21,23,42^ providing an estimate for this parameter, chirality is more difficult to specify from experimental data. Chirality is a critical parameter for tube formation, as these structures are formed by rolling up a flat, honeycomb-like sheet of PduJ hexamers;^23,25^ however, geometric constraints imposed by this hexameric lattice mean that not all diameters are structurally feasible. ^43,44^ Taking these constraints into account, we identified three potential diameters that are consistent with experimental TEM observations—16 nm, 20 nm, and 24 nm—and modeled each with two distinct chiralities: armchair and zigzag (SI Table 2). Armchair nanotubes display high rotational and mirror symmetry, with edges aligned in a smooth, chair-like pattern, while zigzag nanotubes exhibit lower symmetry, with edges arranged in a repeating angular pattern. See Materials and Methods for a detailed description of these calculations.

Henceforth, for clarity, we refer to each tube by its diameter (*e.g.*, “24 nm tube”). All the tubes were assembled with the concave side of the hexamers facing outward. We also performed simulations of convex-outward PduJ tubes (see SI Figure 5) over microsecond timescales. The results indicate that although both orientations yield similar short-timescale dynamics, the concave-outward configuration better reflects the hexamer’s geometry, engages stronger N-terminal interactions, and is likely more stable over longer timescales.

Our simulations show that chirality and radius significantly influence the stability and flexibility of PduJ tubes. Figure 3b shows the trajectory of the tube cross-section for each simulated system. These cross-section traces were generated by connecting the centers of mass of adjacent hexamers using periodic cubic spline interpolation to form smooth, closed curves (see SI Figure 6 for more details). For the PduJ tubes in armchair configurations, the 20 nm tube underwent complete rupture after just 120 ns and failed to maintain an intact tubular structure for the duration of the simulation. In contrast, both the 16 nm and 24 nm armchair tubes remained closed throughout the entire simulation, although their stability varied. The 24 nm tube exhibited the highest structural integrity, while the 16 nm tube displayed slightly reduced stability, as reflected by higher RMSD values throughout the simulation in Figure 3d. In the zigzag configurations, all tubes successfully retained a closed shape throughout the simulation. However, relative to the 24 nm armchair tube, zigzag tubes exhibited greater fluctuations in their circular cross-sectional geometry suggesting that this conformation allows increased flexibility of this circular cross section.

**Figure 3:**
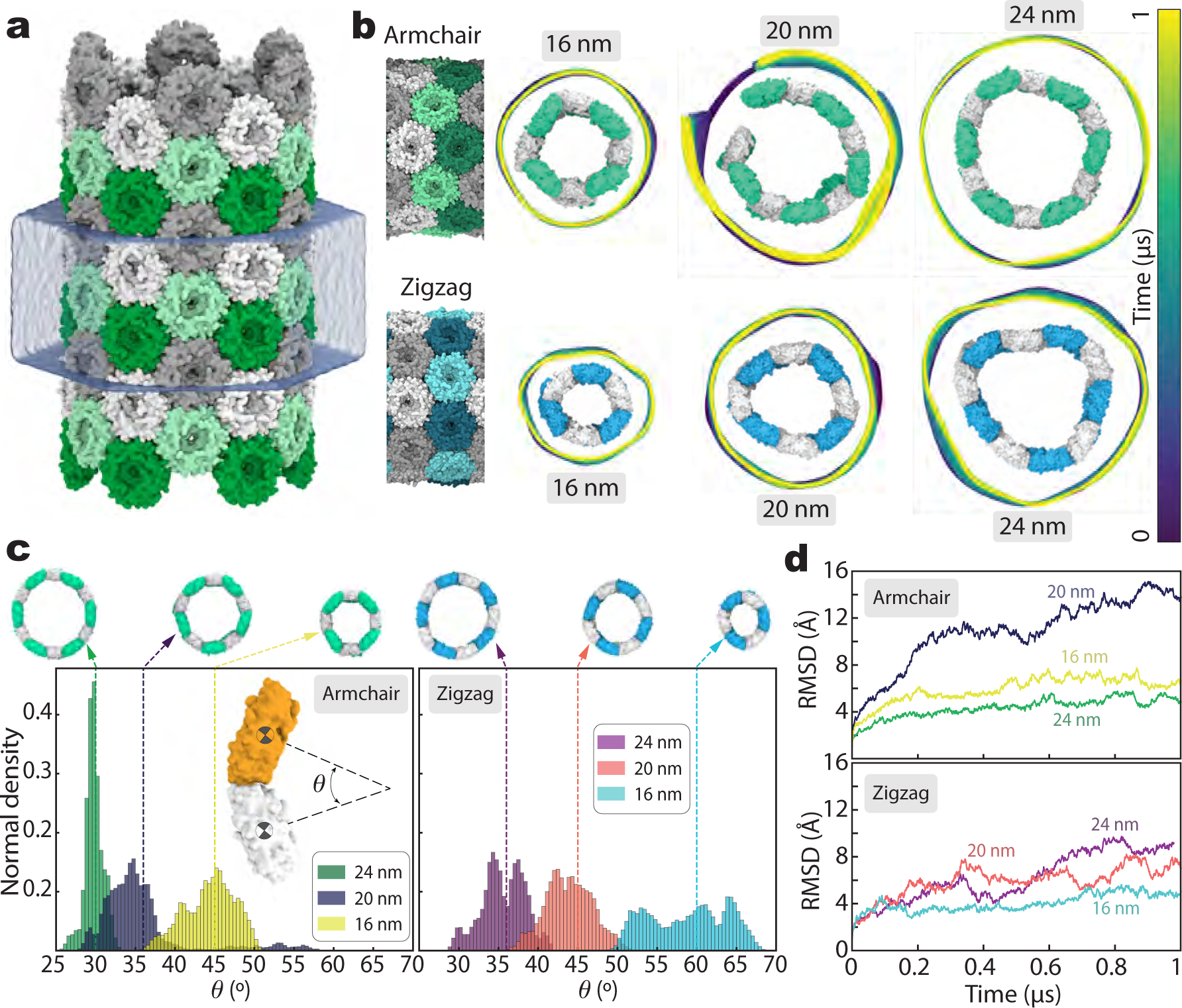
*In silico* structure of a PduJ nanotube. **a,** All-atom model of a PduJ tube with armchair chirality and a diameter of 24 nm, containing 1.5 million atoms. Adjacent rings of the tube are highlighted in green and gray, with alternating hexamers within each ring shown in dark and light shades. The tube is solvated in a hexagonal water box (transparent surface), which serves as the periodic cell. Periodic boundary conditions render the tube effectively infinite. **b,** Trajectories of the tube cross-sections over a 1 *µ*s simulation of six tube systems varying in diameter (16, 20, and 24 nm) and chirality (armchair and zigzag). Cross-section traces are generated by connecting centers of mass of adjacent hexamers using periodic cubic spline interpolation, forming smooth, closed curves. Insets show the final configuration of each tube after the 1 *µ*s simulation. Armchair tubes are shown in green/white, and zigzag tubes in blue/white. **c,** Distribution of bending angles between adjacent hexamers in the armchair (left plot) and zigzag (right plot) simulated systems. The bending angle, *θ*, is defined as the angle between lines connecting each hexamer’s center of mass to the tube center (top right inset). The dashed line indicates the ideal bending angle for a perfectly circular tube, calculated from its radius and number of constituent hexamers. **d,** Tube Root Mean Square Deviation (RMSD) as a function of time. RMSD is calculated for the backbone atoms of each tube relative to their initial configuration.

To more quantitatively interrogate tube stability and flexibility, we measured the hexamer–hexamer bending angle averaged over all frames in the simulation. The bending angle is defined as the angle formed between lines connecting the centers of mass of adjacent hexamers to the central axis of the tube. Figure 3c shows the distribution of the bending angles observed for each system over the course of the simulation. A distribution more sharply centered around the ideal bending angle of the corresponding tube, here, is taken to indicate greater structural stability of a given tube geometry. The 24 nm armchair tube exhibited the sharpest bending angle distribution, indicating a strong preference to preserve the circular shape of the tube structure. Interestingly, this sharpest angle, ∼ 30°, is very close to the minimum angle found in the all-atom simulations of the PMF versus bending angle for PduJ (see SI Figure 4), highlighting the relevance of these bending angle simulations. Conversely, the 20 nm armchair tube, which ruptured during the simulation, showed a broad and irregular bending angle distribution, reflecting substantial structural deformation. In zigzag configurations, the bending angle distribution appeared less sensitive to tube radius; however, all zigzag tubes showed broader distributions than the 24 nm armchair tube, consistent with a higher degree of flexibility observed in our cross section traces. The structural integrity of the tubes was further evaluated by calculating the RMSD of each tube relative to its initial configuration, plotted in (Figure 3d). Over the course of the 1 *µ*s molecular dynamics simulations, the 24 nm armchair tube showed the lowest RMSD (∼4 ^°^A), consistent with high structural stability inferred by cross section traces and bending angle distribution quantification. In contrast, the 20 nm armchair tube displayed a steadily increasing RMSD, reaching values up to ∼20 ^°^A, in line with its observed rupture. Interestingly, the effect of tube diameter on RMSD was less pronounced in the zigzag configuration. Across all tubes with hexamers in zigzag conformation, RMSD values remained relatively stable in the range of ∼ 4–8 ^°^A. Among them, the smallest tube (16 nm) demonstrated the highest stability, which here is indicated by the lowest overall RMSD over the course of the simulation. These findings suggest that while armchair tube stability is strongly dependent on diameter, zigzag tubes are more resilient to size changes, likely due to differences in the geometric or mechanical properties that arise from their differing chiralities. We also characterized tube diameter fluctuations by fitting a circle to the centers of mass of all hexamers projected onto a plane orthogonal to the tube axis (see SI Figure 7a). The distribution of tube diameter for each simulated system, shown in SI Figure 7b, captures how the cross-sectional geometry evolves over time. These fluctuations correlate with the overall structural deviations, as indicated by RMSD traces in Figure 3d.

**Figure 4:**
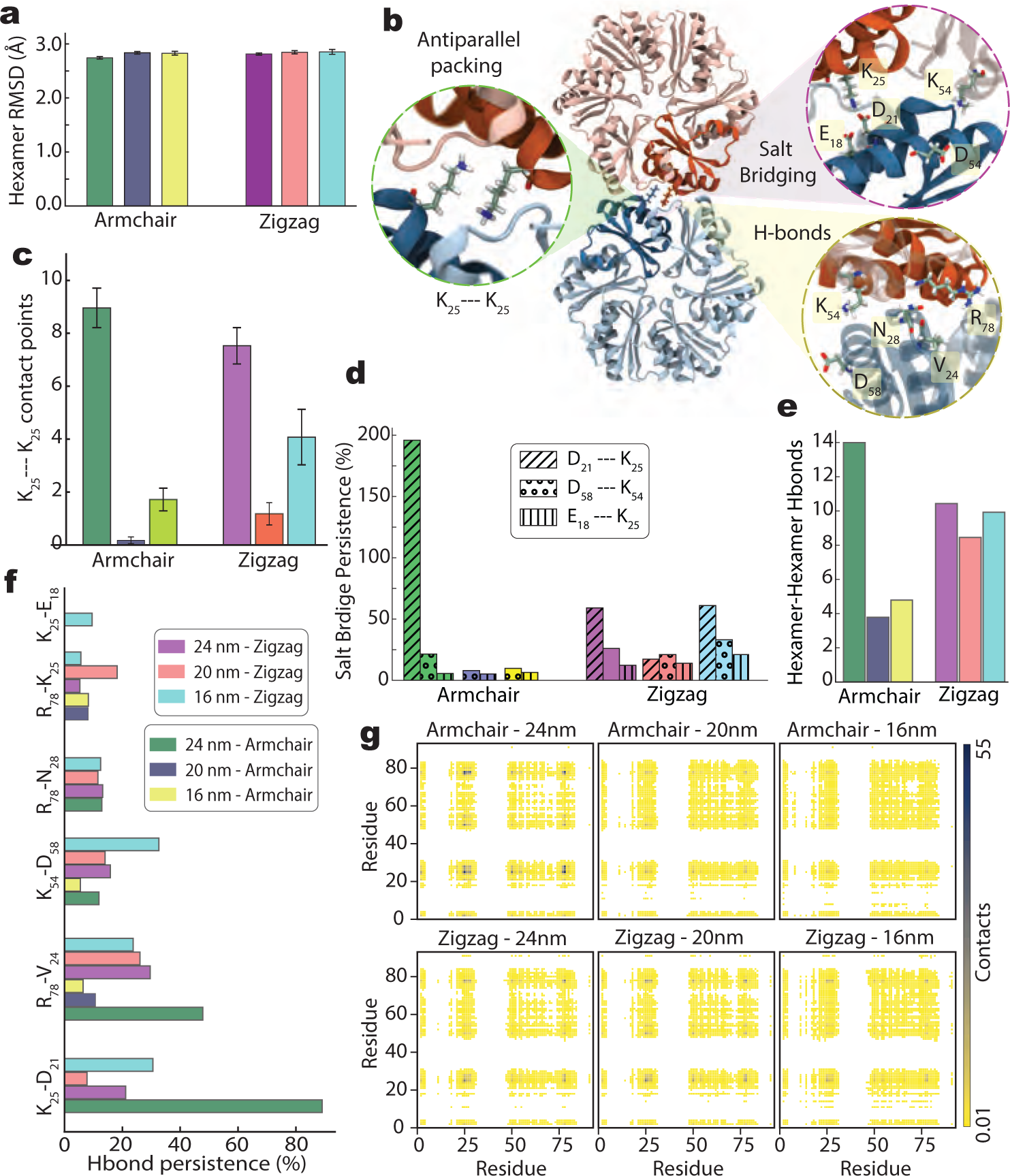
Molecular interactions at the interface of adjacent hexamers in a PduJ nanotube. **a,** RMSD of individual hexamers within the PduJ nanotube, averaged over a 1 *µ*s trajectory. Error bars represent the standard deviation across all constituent hexamers. RMSD values are computed relative to the crystal structure of the PduJ hexamer (PDB ID: 5D6V). **b,** Non-bonded interactions between adjacent hexamers. Zoomed-in insets highlight antiparallel K_25_–K_25_ packing, salt bridges, and hydrogen bonds at the hexamer-hexamer interface. **c-e,** Quantification of inter-hexamer interactions derived from nanotube simulations: (c) average number of K_25_–K_25_ contacts per interface, (d) average salt bridge persistence per monomer, and (e) average number of inter-hexamer hydrogen bonds per hexamer. **f,** Persistence of hydrogen bonds between adjacent hexamers, expressed as the percentage of simulation time during which each bond is maintained. Consistent coloring is used across all panels to represent the same systems; the legend is shown in panel g. All data are averaged over a 1 *µ*s molecular dynamics trajectory. **g,** residue-based 2D maps of hexamer-hexamer contacts extracted from the tube simulations.

This work only includes simulations of armchair and zigzag configurations, where the primary difference between these and chiral configurations lies in the bending angle adopted between hexamers in the final tube structure. By simulating tubes with varying radii in these two conformations, we effectively sampled a broad range of bending angles (Figure 3c), which enables us to estimate the relative stability of other potential chiral structures. If any alternative chiral structure forms with a bending angle of ∼ 30°, which is the bending angle adopted by our most stable 24 nm, armchair configuration tubes, for example, we would expect this alternative chiral structure to be energetically stable. This hypothesis is further supported by our analysis of the interaction energy between two hexamers, where we measured the Rosetta score for hexamer-hexamer bonding as a function of bending angle and separation distance (SI Figure 8). Interestingly, we found that the Rosetta score exhibits a minimum at approximately ∼ 30°, consistent with the most stable tube (24 nm armchair), which shares this same bending angle. Another important consideration is the mechanism of tube formation. If tubes form via the nucleation and merging of small rings, armchair and zigzag configurations are more favorable because they offer higher probability of alignment and successful merging. In contrast, chiral configurations can only merge in a specific orientation, making them less likely to form stable tubes.

To identify the factors underlying the stability variations of tubes with different diameters and chiralities, we first analyzed the RMSD of each hexamer within the tube relative to their coordinates from the crystal structure (PDB ID: 5D6V), Figure 4a and SI Figure 9. Across all simulated systems, the hexamers exhibited an average RMSD of approximately 2.7 ^°^A with a standard error of 0.1 ^°^A, indicating that neither diameter nor chirality significantly impacted the hexamer conformation. This is further confirmed by simulations of a freely equilibrated PduJ hexamer in solution, which has an RMSD of 2.6 ^°^A from the hexamer crystal structure (Figure 1e), suggesting that incorporation into the tubular structure does not affect the conformation adopted by the PduJ hexamers for any of the tube diameters or geometries studied.

Next, we examined the interactions at the hexamer-hexamer junctions that should govern the overall stability of the tube. Previous studies have shown that hydrogen bonding between antiparallel lysine residues (K_25_, Figure 4b) in PduJ hexamers is critical for PduJ assembly, functioning as a sliding lock that stabilizes adjacent hexamers. ^21,45^ To assess the role of this interaction in our systems, we quantified the number of atomic contacts between each K_25_–K_25_ pair for a given tube geometry. A strong correlation emerged between tube stability and the atomic contacts within these antiparallel pairings. Notably, the 24 nm tubes exhibited the highest number of K_25_–K_25_ contacts—8.9 and 7.5 contacts per pair in the armchair and zigzag configurations, respectively (Figure 4c and SI Figure 10). In contrast, the unstable 20 nm armchair tube, which eventually ruptured, displayed a dramatically reduced number of contacts, averaging only 0.1 atomic contacts per pair.

Given that the hexamer–hexamer interface involves several charged residues, including K_25_, D_21_, E_18_, D_58_, and K_54_ (Figure 4b), we next examined the role of salt bridges between oppositely charged amino acids in stabilizing these junctions (Figure 4d). A strong correlation is observed between salt bridge formation and tube stability, particularly in the armchair configuration. Notably, E_18_ and K_25_ form highly persistent salt bridges, with a combined persistence of 196% meaning PduJ monomers form two stable salt bridges between these residues on average, often with neighboring monomers or within itself. This elevated value arises from how salt bridges are defined in the CHARMM force field: the two oxygen atoms in the glutamate side chain carry identical partial charges, and salt bridges are detected based on the proximity between oxygen atoms of acidic residues and nitrogen atoms of basic residues. Consequently, E_18_ can simultaneously engage both of its carboxylate oxygens with the amine group of K_25_, resulting in the identification of two distinct salt bridges and a cumulative persistence exceeding 100%. In the unstable 20 nm and 16 nm armchair tubes, E_18_–K_25_ salt bridge persistence dropped below 10%. In contrast, the flexible zigzag configurations, which exhibited reasonable stability across diameters, maintained an average salt bridge persistence of approximately 25% across different diameters. In armchair tubes, where precise interfacial alignment is essential, even minor deviations in the diameter of the tube can disrupt key interactions and compromise structural stability. Together, these findings suggest that salt bridge formation at the hexamer–hexamer interface is strongly modulated by tube curvature and packing geometry.

A similar correlation is observed between tube stability and hydrogen bonding at the hexamer–hexamer junctions (Figure 4e). In the 24 nm armchair tube—the most stable system—the highest average number of hydrogen bonds (13.9 per hexamer) was measured. In contrast, the two less stable armchair systems exhibited a nearly threefold reduction in hexamer-hexamer hydrogen bonding (3.9 and 5.5 per hexamer, respectively). Interestingly, in the zigzag configuration, hydrogen bonding appeared less sensitive to tube diameter, with all three zigzag systems showing a consistent average of 9.6 ± 1.0 hydrogen bonds. Further analysis of the hexamer-hexamer interface reveals additional key residues in maintaining this interaction, including D_21_, V_24_, and N_28_ (Figure 4b,g). Notably, in addition to the hydrogen bonds it forms with the K_25_ residue on an adjacent hexamer, K_25_ also forms hydrogen bonds with D_21_, with this hydrogen bond reaching a maximum persistence of 85% in the 24 nm armchair system. In agreement with prior experimental mutagenesis studies, R_78_ also emerged as a key residue at the interface,^28,39^ engaging in interactions with V_24_, K_25_, and N_28_, supporting the role of R_78_ in stabilizing hexamer-hexamer interfaces. This observation is consistent with the evolutionary conservation of these specific residues across distant shell proteins, including the metabolosome shell of the myxobacterium *Haliangium ochraceum* and the carboxysome shell of *Halothiobacillus neapolitanus*.^31^ These results indicate that as the bending angle between adjacent hexamers varies with tube radius and chirality, the interfacial residues brought into contact also changes. Consequently, the persistence and geometry of hydrogen bonding at the junctions are modulated by tube architecture, highlighting the sensitivity of these stabilizing interactions to the overall tube curvature.

Further analysis of tube MD trajectories provided two-dimensional contact maps of residue–residue interactions at the hexamer–hexamer interface (Figure 4e). These maps revealed how tube radius influences the nature of interfacial contacts. As the radius decreases, the wedge-like shape of the hexamers causes the subunits to tilt more steeply, increasing the total number of residue–residue contact points between adjacent hexamers. However, this increase is largely driven by transient and low-persistence contacts, while stable, high-persistence interactions (darker regions on the heat maps) are progressively lost. Similarly, with decreasing tube radius, more residues become involved in the interface, but they contribute weakly and inconsistently to the overall stabilization. This analysis highlights that it is not the quantity of contacting residues that determines tube stability, but rather the persistence and strength of specific interactions.

To compare the structural dynamics of PduJ to PduA nanotubes, we constructed and simulated two additional systems comprising 24 nm-diameter PduA tubes with distinct chiralities: zigzag and armchair (SI Figure 11a). This diameter is consistent with experimentally reported PduA tube radii.^21,23,42^ Both zigzag and armchair configurations remained stable throughout the simulations (SI Figure 11b), consistent with the behavior observed for PduJ tubes. Despite the ability of PduA to form stable tubular assemblies, PduJ exhibited a greater propensity to maintain tubular architecture under comparable conditions, in agreement with our cell-free and *in vitro* observations (see the supplementary section S2 for more details).

To better understand the cumulative role of electrostatics in dictating PduA and PduJ assembly, we computed the cohesive energy (non-bonded) of both 24 nm diameter PduJ and PduA tubes in the armchair and zigzag configurations (SI Figure 12). Interestingly, arm-chair PduJ tubes show a significantly higher contribution from electrostatic energy (∼1200 kcal/mol) compared to the energy from Van der Waals interactions (∼150 kcal/mol), highlighting the importance of electrostatic interactions in stabilizing PduJ tubes. Excitingly, the electrostatic contribution for PduJ armchair tubes is significantly higher than in the PduA (∼400 kcal/mol for electrostatic and ∼100 kcal/mol for van der Waals), consistent with our hypothesis that electrostatic interactions stabilize tube formation. The zigzag structures in both WT PduJ and PduA have significantly lower electrostatic contributions to their cohesive energy than the armchair structures.

### Charge-based mutations affect assembly of PduA and PduJ in the native system

The experiments and MD simulations described so far were carried out in non-native environments. However, for applications in metabolic engineering, bioproduction, and antibiotic development, it is necessary to understand how mutated PduA and PduJ assemble within their native context, which involves coassembly with other shell proteins and core enzymes in the native host, *S. enterica*. To assess how charge mutations to PduA and PduJ affect overall MCP nanostructure formation in the native *S. enterica* host, we evaluated MCP assembly upon incorporation of each mutant in the *pdu* operon of the LT2 strain. The assay used here relies on a green fluorescent protein (GFP) tagged with the PduD signal sequence (ssD) that directs the fluorescent GFP reporter to the MCP lumen, thereby allowing visualization of MCP distribution within the cell via fluorescence microscopy.^46^ As expected, cells with WT PduA and PduJ show the presence of two to six small bright fluorescent spots known as puncta, which are assumed to be well-formed MCPs, consistent with prior reports.^21,46^ Other phenotypes that can arise in this assay include elongated stripes of fluorescence corresponding to tube-shaped MCPs, which indicate a shift toward extended assemblies that can form as a result of disrupted vertex capping by the pentameric shell protein PduN.^41^ Alternatively, protein aggregates or polar bodies that appear as larger fluorescent foci at the cell poles indicate disrupted MCP shell formation (Figure 5a). Together, knowledge of these different potential morphologies allows us to interrogate how charge-based mutations disrupt native MCP formation.

**Figure 5:**
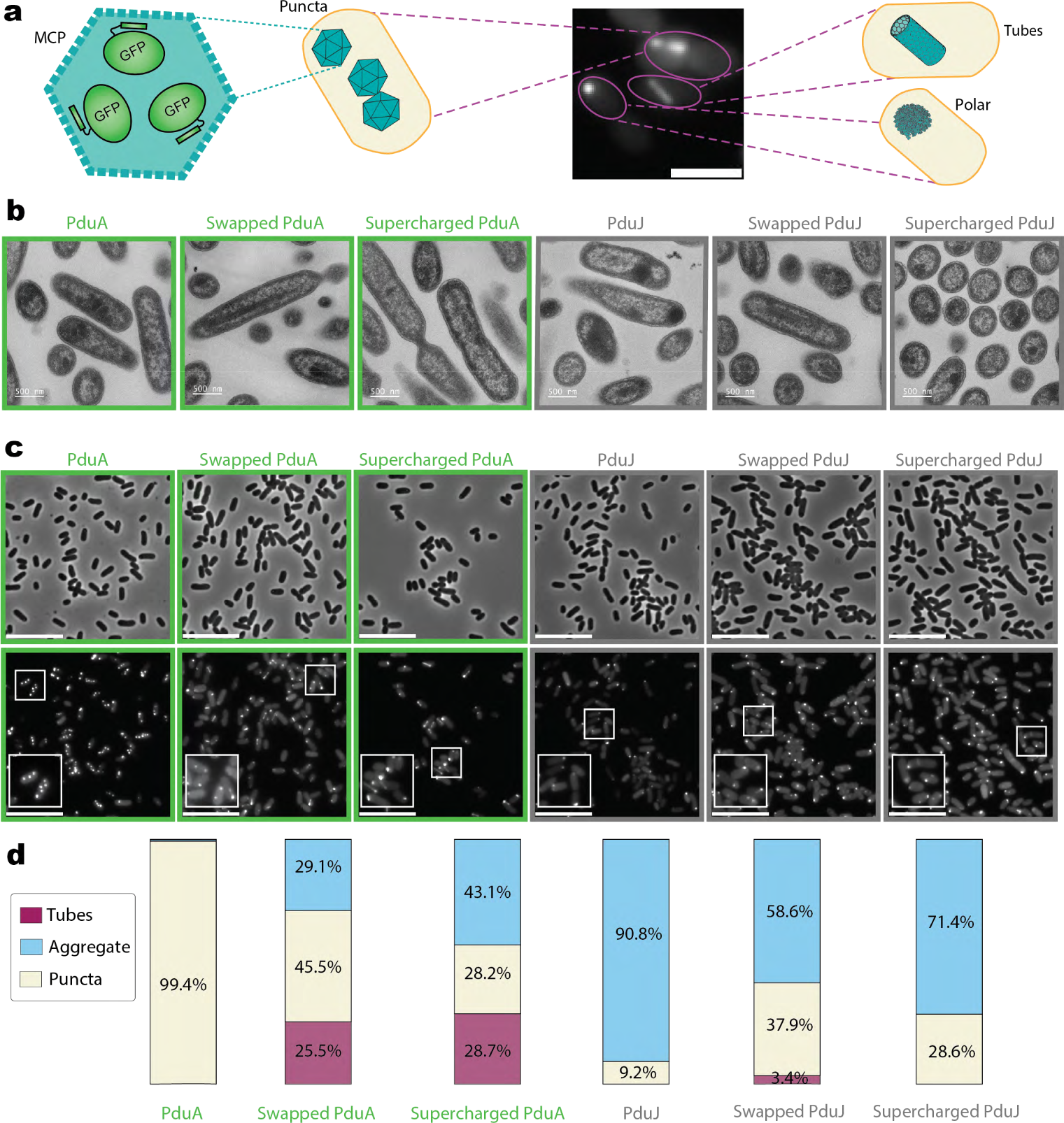
Charge-based mutation effects in LT2 native system. **a,** GFP fluorescence microscopy image of LT2 cells capable of forming MCPs (right), which encapsulate ssD-GFP and produce bright fluorescent puncta distributed throughout the cytoplasm. In addition, LT2 mutant cells can form elongated tubular structures (bottom left) within the cytoplasm, as well as protein aggregates that localize to the cell poles, forming distinct polar bodies (top left). Scale bars = 10 *µ*m. **b,** TEM images of LT2 cell sections reveal distinct structural outcomes. WT PduA forms well-defined MCPs characterized by irregular polyhedral shapes with clear boundaries. In contrast, all PduA mutants exhibit novel tubular structures within LT2 cells. WT PduJ predominantly forms protein aggregates, whereas MCP-like polyhedral structures are recovered in the PduJ mutant strains. Scale bar = 500 nm. **c,** Phase contrast (top row) and GFP (bottom row) fluorescence microscopy images of strains encapsulating ssD-GFP. Small boxes indicate the magnified areas in the inset. Scale bars = 10 *µ*m. **d,** Proportion of structural phenotypes observed in each strain. Data for each strain was collected from three biological replicates (*>*50 cells counted per strain per replicate).

Previous studies suggest that PduA and PduJ play a redundant role in promoting *pdu* MCP assembly, as evidenced by the fact that individual knockouts of either gene do not disrupt MCP assembly, while knockouts of both ablate MCP formation entirely.^21,47^ Because PduA and PduJ exhibit this partial functional redundancy in the native *Salmonella* system, isolating the impact of a given PduA or PduJ mutant requires testing each mutant in a strain where WT PduA or PduJ is knocked out. To this end, all PduA and PduJ variants (WT, swapped, supercharged) expressed off the *pduA* locus in a strain with the WT *pduJ* gene knocked out, such that only the protein encoded at the *pduA* locus would be able to interact with the remaining shell proteins and contribute to the formation of the irregular polyhedral MCP (Figure 5c).

Collectively, study of charge variants in the native system suggests that both hexamer charge and expression locus affect the MCP structures formed. Using the fluorescence microscopy assay described above, we find that 99.4% of the cells expressing WT PduA at the *pduA* locus with *pduJ* knocked out contained three or more flourescent puncta, indicative of proper MCP assembly^21^ (Figure 5d). In contrast, only 9.2% of cells expressing WT PduJ from the *pduA* locus contained three or more fluorescent puncta, and instead primarily contained protein aggregates and diffuse GFP fluorescence, indicating impaired MCP assembly, possibly due to differences in expression from the non-native locus. The strain with swapped PduA produced significantly more cells with three or more puncta (45.5%) than the WT PduJ strain (*p <* 0.001) but fewer than the WT PduA strain (*p <* 0.001). The swapped PduJ strain also produced significantly more cells containing three or more puncta (37.9%) than the WT PduJ strain (*p <* 0.001) indicating that hexamer charge can mediate MCP assembly. The strain harboring supercharged PduJ produced both cells with normal MCPs (28.6%) and cells with protein aggregates (71.4%). Interestingly, while strains with supercharged PduJ showed comparable levels of MCP formation, swapped PduJ conferred a modest increase in assembly relative to WT (Figure 5c,d). This suggests that swapped and supercharged PduJ variants either more closely mimic PduA in their ability to stimulate MCP assembly, or that the expression defect typically associated with heterologous expression of *pduJ* at the *pduA* locus is less pronounced for these variants.

Tubular MCP structures were observed alongside normal MCPs and protein aggregates in some variants tested. Specifically, tubular MCP structures were detected in 25.5% of cells with swapped PduA, 28.2% of cells with supercharged PduA, and 3.4% of cells with swapped PduJ, whereas cells with supercharged PduJ contained no detectable tubular structures (Figure 5d). The presence of tubular shells for some of these hexamer variants suggests that, although these hexamer variants are still able to interact with some of the other hexagonal shell proteins (such as PduB, which is responsible for linking the MCP core and shell ^48^), some fraction of these variants disrupt pentamer (PduN) incorporation into the shell, which has been shown to mediate the formation of tubular MCPs.^41,49^

To confirm the results of our fluorescence microscopy assay, we also visualized the subcellular structures formed by the expression of the *pdu* operon with our various hexamer variants using TEM imaging of sectioned and stained cells. These TEM images revealed structures consistent with our fluorescence microscopy analyses across all strains (Figure 5b). TEM on cells expressing WT PduA showed the expected distinguished, flat, angular boundaries characteristic of Pdu MCPs. We note that because Pdu MCPs are irregular and polyhedral, the observed structures associated with MCPs in these thin cell sections are morphologically heterogeneous, but can be distinguished by their defined boundaries. In contrast to cells expressing WT PduA with other compartment components, cells expressing WT PduJ overwhelmingly showed aggregates at or near the cell pole with no distinct boundaries, consistent with the protein aggregates observed by fluorescence microscopy for this strain. Both swapped and supercharged PduA exhibit a combination of MCPs, aggregates, and long parallel tubes, the latter of which appear with defined, stripe-like boundaries that correspond to the walls of protein tubes with high protein content between them observed in thin cell section TEM of strains with the PduN pentamer knocked out^41^ (Figure 5d). Similar to the smaller tubes formed by PduA and PduJ alone, these tubular MCPs packed tightly together into bundles within the LT2 cytoplasm and are typically oriented along the longitudinal axis of the cell. Swapped PduJ cells have some tubes but, like supercharged PduJ, yielded primarily MCPs and polar aggregates.

The observation of tubular structures in strains expressing PduA variants indicates that increasing the net charge of the hexamer disrupts the balance between shell closure and extended growth, thereby favoring the formation of tubular assemblies and protein aggregates over closed polyhedral MCPs. A plausible explanation for this observation is that charge mutations modify preferred interaction angles between hexamers while preserving coplanar hexamer-hexamer contacts, thereby preventing pentamers from priming the shell into a closed, irregular polyhedral architecture.^41^ This suggests that the PduA lattice is highly sensitive to charge perturbations and that such perturbations can easily tip the balance of this lattice toward elongated assemblies and aggregates over closed MCPs. In contrast, charge modulation in PduJ produced the opposite effect in the context of MCP assembly. Although WT PduJ expressed at the *pduA* locus did not result in well-formed polyhedral MCPs, altering its surface charge enabled some degree of proper shell closure, suggesting that the mutations reactivated or stabilized a latent capacity for the formation of closed MCPs. Such charge changes may enhance pentamer-hexamer interactions or strengthen key bent junctions required for MCP closure, thereby favoring assembly of a closed, polyhedral MCP. Taken together, this suggests that PduA is more dependent on precise electrostatic tuning for achieving balance between closure and extension, whereas PduJ is more tolerant to charge perturbations in this context.

Although supercharged PduJ and supercharged PduA share the same overall net charge (−12 e), they assemble into different structures depending on the context. Under native conditions, which require coassembly with other MCP shell components, supercharged PduJ forms a mixture of MCPs and aggregates, whereas supercharged PduA forms tubes, MCPs and aggregates (Figure 5d). When expressed alone in CFPS, supercharged PduJ assembles predominantly into large fibrilar structures, while supercharged PduA forms a mixture of large tubes and sheet-like tubular bundles (Figure 2). These results indicate that the net charge alone does not determine the surface electrostatic landscape or assembly behavior of these hexameric proteins. This is likely because assembly pathways are not governed by charge alone, but also by cooperativity with other shell proteins, which influence factors like hydrogen-bonding capacity and the propensity to form salt bridges.

## Conclusions

By integrating heterologous and cell-free expression studies with native co-assembly experiments, we demonstrate that charge-altering variants provide a systematic framework for modulating the morphology of hexameric proteins within Pdu bacterial microcompartments (MCPs). Molecular dynamics simulations elucidate the molecular-level interactions governing these assemblies. Specifically, in tubular structures, our results reveal a robust correlation between hydrogen bonding, salt-bridge persistence, and overall stability, highlighting the role of curvature in modulating these stabilizing interactions.

We show that protein-specific geometries (*e.g.* tubes and sheets) can be modified by analyzing the key interactions in the different hexameric proteins. Wild type (WT) PduJ, which has higher overall negative charge than WT PduA, favors the formation of extended tubular structures in cell-free environments. Molecular dynamics simulations demonstrated that precise interfacial alignment is essential and even minor deviations in the diameter of the tube can disrupt key interactions and compromise structural stability. Our findings suggest that salt bridge formation at the hexamer–hexamer interface is strongly modulated by tube curvature and packing geometry. In PduJ, specific inter-subunit contacts or geometric constraints promote curvature and disfavor flat hexameric lattices. While PduA tubes are capable of forming stable assemblies, PduJ exhibits a higher propensity for maintaining tubular architecture under similar conditions. Electrostatic interactions dominate the cohesive energy of the hexamers in WT PduJ tubes much more than in WT PduA tubes, demonstrating that increased electrostatic interactions favor tube formation.

We find that even when hexameric variants share the same overall net charge, they adopt distinct structural configurations, indicating that assembly cannot be explained by charge alone. Instead, hydrogen-bonding patterns and salt-bridge networks play a critical role. Consistent with this, swapped variants did not produce symmetrical morphological changes. Since there is ∼20% difference in sequence of amino acids between PduA and PduJ and the local orientation and dynamics of water molecules near a protein surface, which dictate the assembly route, depend on the polarity of the amino acids,^50^ we cannot expect electrostatics to be alone responsible the resulting morphologies. Increasing the negative charge, however, promotes the formation of tubular structures, consistent with electrostatic interactions effects, which favors one dimensional growth in strongly charged self-assembling units.^40,51^ In native systems, WT PduA appears intrinsically optimized for polyhedral closure, while all PduA mutants (which have increased overall charge) exhibit tubular structures. In contrast, WT PduJ predominantly forms protein aggregates, whereas MCP-like polyhedral structures are recovered by PduJ mutants with charge variation. Our results support that charge-altering mutations can redirect assembled morphologies by tuning electrostatic balance, revealing alternative assembly routes that emerge under specific charge-patterning conditions. Our heterologous overexpression and cell-free expression experiments show that these platforms provide powerful and highly controllable environments for probing and directing self-assembly. The ability to program distinct morphologies through specification of the assembling protein subunit holds significant promise for achieving high mechanical performance and expanding the functional versatility of protein-based materials. Together, these insights deepen our understanding of how subtle electrostatic and geometric factors govern self-assembly and lay the groundwork for designing and controlling protein-based nanostructures.

## Methods

### Experimental Methods

#### Plasmid and strain construction

All mutated variants of *pduA* and *pduJ* were ordered as gBlocks from Twist Biosciences (SI Table 3) and PCR-amplified using primers encoding Golden Gate–compatible overhangs (see Table 4). Amplified gene fragments were then cloned into either a Golden Gate–compatible pBAD33t vector containing a *p15A* origin of replication and a chloramphenicol resistance cassette, or into a pJL1 parent vector, which includes an IPTG-inducible T7 promoter, a kanamycin selection cassette, and a ColE1 origin of replication for cell-free protein synthesis experiments. A C-terminal FLAG tag was included on all plasmid-encoded PduA and PduJ charge variants. All cloning was performed using *Escherichia coli* DH10b cells.

The *pdu* operon of *Salmonella enterica* serovar Typhimurium LT2 was modified using the *λ* Red recombineering method.^52^ Genetic modifications were first performed to replace the gene at the locus of interest with a *cat/sacB* selection marker amplified from the TUC01 genome using primers containing 5^1^ and 3^1^ homology to the locus. The PCR products were used to incorporate the triple point charge *pduA* and *pduJ* variants at the *pduA* locus with the *pduJ* gene knocked out,^21^ to create the corresponding charge variant strains. These strains were amplified from their respective plasmid using primers with 41-51 base pairs of homology to the C-terminal region of the *pduA* locus. We confirmed that the locus had been correctly modified by amplifying the region with PCR and checking its sequence using Sanger sequencing.

#### Growth conditions for linked cell assay

The linked cell assay to test self-assembly of individual hexamer variants upon heterologous overexpression *in vivo* was carried out as previously described.^21^ All variants were cloned into pBAD33t expression vectors with a C-terminal FLAG tag as described above. Overnight cultures were inoculated from single colonies in 5 mL of lysogeny broth—Miller formulation (LB-M) supplemented with 34 *µ*g/mL chloramphenicol for plasmid maintenance, and grown for 15-16 hours at 37 *^◦^*C with shaking at 225 rpm in 24-well blocks. These saturated overnight cultures were then subcultured 1:100 into 5 mL of fresh LB-M, again supplemented with 34 *µ*g/mL chloramphenicol in 24-well blocks. These new cultures were grown for 90 minutes (or to an OD_600_ of 0.2-0.5) at 37 *^◦^*C with shaking at 225 rpm and were then induced via addition of 50 *µ*L of 20 wt% L-(+)-arabinose to a final arabinose concentration of 0.2 wt%. After induction, cultures were grown for 4-6 hours to allow the linked cell phenotype to develop prior to imaging using phase contrast microscopy (see below).

#### *Salmonella* growth conditions for GFP encapsulation assay

Overnight cultures of recombineered LT2 strains shown in table 1 were transformed with the pBAD33t-ssD-GFPmut2 reporter plasmid (CMJ069). They were grown from a single colony into fresh 5 mL LB-M supplemented with 34 *µ*g/mL chloramphenicol grown at 37 *^◦^*C with 225 RPM shaking in 24-well blocks for 15–16 hours. The overnight cultures were then subcultured 1:500 (10 *µ*L) into 5 mL of of fresh LB-M with 0.02% (*w/v*, final concentration) L-(+)-arabinose, 34 *µ*g/ml chloramphenicol, and 0.4% (*v/v*, final concentration) 1,2-propanediol (1,2-PD) in 24-well blocks. Cells were grown at 37 *^◦^*C with shaking at 225 RPM for at least 6 hours following subculture. After incubation, 700 *µ*L of each culture was transferred to a 1.5 mL microcentrifuge tube and centrifuged at 4,000 × *g* for 1.5 minutes. Following centrifugation, 600 *µ*L of the supernatant was carefully removed, and the cell pellet was resuspended in the remaining 100 *µ*L. Resuspended cells were then imaged using a combination of phase contrast and fluorescence microscopy (see below).

#### Phase contrast and fluorescence microscopy

Phase contrast and fluorescence microscopy was performed using Fisherbrand ^TM^ frosted microscope slides (Thermo Fisher Scientific Cat# 12-550-343) and 22 mm × 22 mm, #1.5 thickness cover slips (VWR Cat# 16004-302). Imaging was performed on a Nikon Eclipse Ni-U upright microscope with a 100X oil immersion objective using an Andor Clara digital camera. Image acquisition was performed using NIS Elements Software (Nikon). GFP fluorescence images were captured with a C-FL Endow GFP HYQ bandpass filter, using a standardized exposure time of 200 ms. See supplementary section S1 for more details on the quantification protocol.

#### Ultra-thin section transmission electron micrscopy

Bacterial cells expressing the pBAD33t-ssD-GFPmut2 reporter were grown under the GFP encapsulation assay conditions, after which 1.5–2 mL of each culture was transferred to microcentrifuge tubes and centrifuged at 3,400 × g for 1 min. The resulting pellets were fixed in a solution containing 2.5% electron microscopy (EM)-grade glutaraldehyde and 2% paraformaldehyde in 0.1 M 1,4-piperazinediethanesulfonic acid (PIPES) buffer. Post-fixation was carried out using 1% osmium tetroxide, followed by dehydration through a graded ethanol series. Samples were then infiltrated and embedded in EMBed 812 epoxy resin, which was cured at 60 *^◦^*C for 48 hours. Ultra-thin sections (approximately 50 nm thick) were cut using a Leica UC7 ultramicrotome equipped with a diamond knife and collected on slotted grids coated with Formvar and carbon. Sections were subsequently stained with 3% uranyl acetate and Reynold’s lead citrate. Imaging was performed on a Hitachi HD2300 STEM operated at 200 kV, using both phase contrast transmission and high-angle annular dark field (HAADF) modes.

#### Cell extract preparation for cell-free protein synthesis (CFPS)

*E. coli* BL21(DE3) (Life Technologies) cells were prepared into cell-free extracts via growth, harvest, lysis, and preparation as previously described, see supplementary section S1 for more details.

#### CFPS Reactions

Cell-free protein synthesis (CFPS) was conducted according to methods described in detail in supplementary section S1.

#### Confocal/super-resolution microscopy of PduA and PduJ variants in CFPS reactions

For imaging the cell-free produced PduA and PduJ assemblies, the CFPS reactions were prepared with a 1:100 dilution of DYKDDDDK tag monoclonal antibody (FG4R), DyLight^TM^ 550 (Invitrogen, Catalog No. MA1-91878-D550) in a *µ*-Slide 18-well glass bottom cell imaging chamber, allowing direct examination with a Nikon SoRa spinning disk confocal microscope equipped with an air 20× (0.8 NA) objective lens and a silicone-immersion 40× objective lens (1.25 NA). For image acquisition, a Yokogawa CSU-W1 dual-disk spinning disk unit with 50 *µ*m pinholes, a SoRa spinning disk, and a Hamamatsu ORCA-Fusion digital CMOS camera were used. Microscope control was performed using Nikon imaging software NIS-Elements AR. All variants were imaged on at least three separate days. The SoRa microscope is housed in the Northwestern University Biological Imaging Facility, supported by the Northwestern University Office for Research.

### All-atom models of PduJ and PduA tubes

The structure of PduJ nanotubes with armchair and zigzag configurations is inspired by terminology commonly used for single-walled carbon nanotubes, which originate from rolling 2D graphene sheets into cylindrical forms, which was similarly adopted by researchers exploring potential morphologies of PduA tubes. ^45^ The nanotube diameter, *d*, can be described in terms of the hexagonal lattice edge length, *a*, and is calculated as follows:

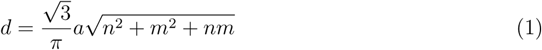

Here, *n* and *m* are the chiral indices that define the wrapping vector of the tube and ultimately determine its geometry and symmetry. In the armchair configuration, both indices are equal (*n* = *m*), whereas in the zigzag configuration, one index is zero (*m* = 0). Based on measurements from the PduJ hexamer model and tube diameters observed in TEM images, both armchair and zigzag conformations were adopted to model PduJ nanotubes, as summarized in SI Table 2.

To build the all-atom model of the tubes, we arranged the hexamers into a ring using either zigzag or armchair chirality at the target radius. To extend the ring into a tube, we duplicated the ring to make a 4-layer tube. To avoid steric clashes between the hexamers, in both ring and tube structures there is at least 3 ^°^A distance between the adjacent hexamers. After centering and aligning the tube along the *z* -axis, it was submerged in a hexagonal water box with 30 ^°^A of water separating it from each side. Next, Na^+^, and Cl*^−^* ions were added to the system to produce a 0.1 M electrolyte solution. The system underwent 4800 steps of energy minimization and NPT equilibration in which the backbone atoms of the protein were harmonically restrained to their initial coordinates using the following parameters. For the first 2 ns, the spring constant of each restraining potential was 1.0 kcal/mol/^°^A^2^, which was then reduced to 0.5, 0.1, and 0.01 kcal/mol/^°^A^2^ in intervals of 3, 4, and 20 ns, respectively. Next, the harmonic potential was removed and the system was equilibrated for 12 ns. Finally, for the production runs, we cut the first and last rings including any solution around them from the equilibrated tubes, and kept the two middle rings. Through periodic boundary conditions (PBC), the tubes were effectively infinite. The height of the simulation box was set to the *z* -distance between maximum and minimum of the two middle rings, plus the equilibrated gap between the rings. As result of removing the solution around the first and fourth rings, additional ions were added if required to maintain a net neutral charge in the system. See Supporting Information for a detailed description of the methods and materials for MD simulations.

## Contributions

A.G., M.O.d.l.C., B.M. and D.T.E. designed the project with input and initial contributions from C.W., N.W.K., and C.E.M.. A.G. and C.E.M performed experiments, with additional contributions from J.M. B.M. performed molecular dynamics simulations of proteins and tube morphologies, with contributions from A.G.. C.W. performed PMF calculations of bending angle of two interacting proteins. A.G., B.M., D.T.E., and M.O.d.l.C. contributed to analysis and interpretation of results and wrote the first draft of the manuscript. N.W.K., C.E.M, and D.T.E. contributed to editing the manuscript.

## Acknowledgments

We thank Eric W. Roth for their assistance with TEM. The work was funded by the National Science Foundation (NSF) via the Northwestern University MRSEC grant number NSF-DMR 2308691. This research was supported in part through the computational resources and staff contributions provided for the Quest high performance computing facility at Northwestern University which is jointly supported by the Office of the Provost, the Office for Research, and Northwestern University Information Technology. The authors thank David Glass, Sai Kumar Ramadugu, and Scott Coughlin for their assistance with implementing NAMD3 on the Quest supercomputer.

