## Supplementary material for "Engineering the Self-assembly of Bacterial Microcompartment Shell Proteins via Charged Mutations": SI

### 1. Details of Experimental Protocols

#### Phase contrast and fluorescence microscopy quantification

Brightness and contrast adjustments were applied uniformly across experiments using ImageJ software.[1] Cell length measurements were collected using the segmented line tool in ImageJ-2 software.[1] Cell counts were performed on images with adjusted brightness and contrast. To quantify the fraction of linked or elongated cells, the cells were categorized on the basis of the number of cleavage furrows per cell. Groups containing three or more cells separated by visible cleavage furrows were classified as linked, while those with two or fewer cells (or with one or no identifiable cleavage furrows) were considered unlinked. Each individual cell body within a linked group was counted as a separate cell event (see Figure 2). To quantify the varying phenotypes present for the GFP encapsulation assay,the presence of two to six small bright fluorescent spots per cell was counted as normal MCP formation. Elongated tubes retained within the cytoplasm and protein aggregates at the cell poles were each counted as separate phenotypes (see Figure 5).

#### Cell extract preparation for cell-free protein synthesis (CFPS)

Cells were prepared as previously described by Karim and Jewett.[2] First, cells were cultured in full-baffled Tunair flasks with 1 L of 2x YTPG (16 g/L tryptone, 10 g/L yeast extract, 5 g/L NaCl, 7 g/L potassium phosphate monobasic, 3 g/L potassium phosphate dibasic, 18 g/L glucose, pH 7.2) at 37 °C and shaking at 225 RPM. T7 RNA polymerase production was induced using 1 mM isopropyl-$\beta$-D-thiogalactopyranoside (IPTG) when cultures reached an $OD_{600}$ of 0.6–0.8. Cells were harvested at $OD_{600}$ of 3.0 $\pm$ 0.1 by centrifugation at 5,000 $\times$ g for 10 minutes at 4 °C. The supernatant was decanted, and the remaining pellets were washed three times with cold S30 buffer (10 mM tris acetate, pH 8.2, 14 mM magnesium acetate, and 60 mM potassium acetate) and subsequently centrifuged at 10,000 × g at 4 °C. Pellets were then flash frozen with liquid nitrogen and stored at -80 °C. Cells were then thawed and lysed using the EmulsiFlex-B15 homogenizer (Avestin) in a single pass at a pressure of 20,000–25,000 psi. Debris was removed by centrifugation (12,000 × g, 30 min, 4 °C). The supernatant from this step was collected as the final cell extract and was divided into 50 $\mu$L aliquots, flash frozen in liquid nitrogen, and stored at -80 °C until use.

#### CFPS Reactions

Cell-free protein synthesis (CFPS) was conducted according to methods described previously described in.[2,3] To produce proteins for analysis by confocal microscopy, 40 $\mu$L CFPS reactions were set up in $\mu$-Slide 18-well high glass bottom chambers (IBIDI, Cat.# 81817). Each CFPS reaction contained: 10 mM $Mg(Glu)_2$, 10 mM $NH_4$(Glu), 130 mM K(Glu), 1.2 mM adenosine triphosphate (ATP), 0.85 mM guanosine triphosphate (GTP), 0.85 mM uridine 5′-triphosphate (UTP), 0.85 mM cytidine 5′-triphosphate (CTP), 0.034 mg/mL folinic acid, 0.171 mg/mL tRNA, 33.33 mM phosphoenol pyruvate (PEP), 2 mM of each of the 20 standard amino acids, 0.33 mM $NAD^+$, 0.27 mM CoA, 4 mM coenzyme A (CoA), 1 mM putrescine, 1.5 mM spermidine, and 57 mM 4-(2-hydroxyethyl)-1-piperazineethanesulfonic acid (HEPES), 0.3 volume fraction *E. coli* BL21(DE3) extract, and a 1:100 dilution of DYKDDDDK Tag Monoclonal Antibody (FG4R), DyLight™ 550 (ThermoFisher Scientific, Cat.# MA1-91878-D550). To initiate CFPS reactions, 13.33 ng/$\mu$L pJL1 plasmid subcloned with each C-terminally FLAG-tagged hexamer variant (see above) was added to the reaction and mixed. The CFPS reactions were pipetted into $\mu$-Slide 18-well high glass bottom chambers (IBIDI, Cat.# 81817) and incubated at 30 °C for $>$2 hours to allow protein expression and assembly to occur.

### 2. Details of MD Simulations

#### General molecular dynamics methods

All molecular dynamics (MD) simulations were conducted using NAMD3,[4] with protein and lipid molecules parameterized by the CHARMM36 force field[5,6] and water molecules modeled using TIP3P.[7] Periodic boundary conditions were applied in all three dimensions. Electrostatic interactions were computed using the Particle Mesh Ewald (PME) method on a 1.2 Å grid.[8] Van der Waals interactions were smoothly truncated between 10 and 12 Å. Unless stated otherwise, the Hydrogen Mass Repartitioning (HMR) approach[9] was utilized to allow a 4 fs integration time step, along with the SETTLE[10] and RATTLE[11] algorithms to constrain covalent bonds involving hydrogen atoms in water and protein molecules, respectively. Both short-range and long-range electrostatic interactions were evaluated at every simulation step. In simulations conducted under the canonical ensemble (NPT), a Langevin thermostat was applied to all non-hydrogen atoms, using a damping coefficient of 5 ps$^{-1}$ to regulate the system temperature at 295 K. For simulations in the isothermal-isobaric ensemble (NPT), pressure was maintained at 1 bar using the Nose-Hoover Langevin piston method.[12,13] Energy minimization was carried out using the conjugate gradient algorithm. Visualization and data analysis were performed with the Visual Molecular Dynamics (VMD) software.[14]

#### All-atom models of PduJ and PduA Hexamers

The all-atom hexamer models were constructed using the crystal structures of the PduJ $K_{25}A$ mutant (PDB ID 5D6V[15]) and PduA from *Salmonella enterica* Typhimurium (PDB ID 3NGK[16]). As the PduJ crystal structure contained a mutation at residue 25, the alanine was replaced with lysine to restore the wild-type sequence. Missing atoms in the crystallographic structures and the mutation correction were incorporated using the *psfgen* package in VMD.[14] Each hexamer was subsequently placed in a $14 \times 14 \times 14$ $nm^3$ water box, and NaCl was added to achieve a final concentration of 100 mM. The system was first subjected to 5,000 steps of energy minimization, followed by a 15-ns equilibration phase in which all non-hydrogen protein atoms were restrained to their crystallographic positions. The restraint spring constants were initially set to 5 kcal/mol/$\mathring{A}^2$ and gradually reduced by 0.1 kcal/mol/$\mathring{A}^2$ every 5 ns. A subsequent 20-ns equilibration step was performed with only the protein backbone atoms restrained using a 0.1 kcal/mol/$\mathring{A}^2$ force constant. Finally, each system underwent an unrestrained production simulation lasting 350 ns.

#### Analysis of molecular dynamics trajectories

VMD and MDAnalysis were used for the analysis of simulation trajectories, with VMD also employed for visualization. Root-mean-square deviation (RMSD) and hydrogen bond analyses were performed using the RMSD Tool and HBonds Plugin in VMD. RMSD calculations for the PduJ and PduA hexamers were performed with respect to their corresponding crystal structures (PDB ID: 5D6V[15] and PDB ID: 3NGK,[16] respectively). For tube structures, RMSD was calculated relative to the equilibrated structure, defined as the final frame of the infinite system simulations. Hydrogen bonding between hexamers was assessed using a donor–acceptor distance cutoff of 3.0 Å and an angle cutoff of $30^\circ$. Salt bridge interactions between hexamers were identified using in-house Python and Tcl scripts. A salt bridge was considered to be formed if the distance between any oxygen atom of an acidic residue and any nitrogen atom of a basic residue was within 3.2 Å in at least one simulation frame. Contact points between the hexamers were identified using an interatomic distance cutoff of 4.5 Å.

#### MD simulation of PduA tubes

To compare the structure and dynamics of PduA and PduJ tubes, we constructed two additional systems, each containing a 24 nm-long PduA tube in either a zigzag or armchair configuration. The simulation setup and conditions were identical to those used for the PduJ tube simulations. Our simulations revealed the armchair configuration more effectively preserved the tube's circular cross-sectional geometry (SI Figure 11c). RMSD profiles of both PduA tubes were comparable, with RMSD values stabilizing around ∼8 Å SI Figure 11d, similar to those of the 24 nm zigzag PduJ tube. The most persistent inter-hexamer hydrogen bond was identified between residues $R_{79}$ and $V_{25}$. Notably, the total number of hydrogen bonds between adjacent hexamers in the PduA tubes was reduced relative to PduJ, by factors of 1.5 and 2.1 for the zigzag and armchair configurations, respectively. Although PduA tubes exhibited the same key salt bridge interactions at the hexamer interfaces as observed in PduJ SI Figure 11g, the persistence of these interactions was considerably lower, suggesting weaker salt bridging at the junctions. No significant structural disruptions were observed within individual hexamers for either chirality over the course of the simulations SI Figure 11h. In summary, while PduA tubes are capable of forming stable assemblies, PduJ exhibits a higher propensity for maintaining tubular architecture under similar conditions, consistent with our observations in cell-free and *in vitro* experiments (Figure 2).

### Concave in-out configuration of the PduJ tubes

PduJ hexamers are wedge-shaped when viewed perpendicular to their sixfold symmetry axis and exhibit two distinct faces: a concave (inward-curving) side and a convex (outward-curving) side, Figure 1b. A key unresolved question in the field is whether hexamers assemble into tubes with the concave face oriented outward or inward. Geometrically, the wedge-shaped shape of each hexamer resembles a voussoir, allowing the units to interlock and pack tightly into a stable curved structure, supporting the hypothesis that the concave face is oriented outward in the assembled tube SI Figure 5a. Although both orientations support stable tube formation over microsecond simulations, the concave-outward configuration is consistent with the hexamer's geometry and exposure of functionally important N-terminal regions exhibits stronger interfacial interactions, including more persistent salt bridges and $K_{25}$–$K_{25}$ contacts. These differences suggest that, despite comparable structural dynamics, the concave-outward arrangement is thermodynamically more stable over longer timescales. Together, these results suggest that while both tube configurations exhibit similar structural dynamics over microsecond timescales, the concave-outward arrangement is likely to be more stable over longer timescales.

#### Calculation of the bending potential

We calculate the bending potentials of mean force (PMF) shown in SI Figure 4a, using the same method as in our previous work.[17,18] The bending potential for PduA was previously published, but is included here for comparison. The initial structure for the atomistic model of the hexamer interfaces were generated using the same protocol for PduA/PduA interfaces which was previously described.[17] Additional mutations were made using the Pymol[19] mutagenesis wizard. The surface potentials were also calculated using PyMol. The bending potential, $V(\theta_B)$, between two hexamers, $\text{Hex}_1$ and $\text{Hex}_2$, is calculated from the forces on $\text{Hex}_2$ at a given bending angle, according to the definition of the force, $F_{\theta_B}$:

$$F_{\theta_B} = -\frac{\partial V}{\partial \theta_B} \tag{1}$$

The quantity $\theta_B$ is defined in SI Figure 4b as the dot product of the normal vectors of $\text{Hex}_1$ and $\text{Hex}_2$. We bias the center of mass distance between $\text{Hex}_1$ and $\text{Hex}_2$ in the $z$-direction using a harmonic spring. We also restrain the backbone of $\text{Hex}_1$ in all directions. We then measure the forces in the $z$-direction as logged in the simulation. These forces in the $\theta_B$ direction are the component of the total force in the z-direction in the $\theta_B$ direction:

$$F_{\theta_B} = F_{spring} cos(\theta_B) \tag{2}$$

The other components of the force are canceled by the restraints of $Hex_1$. The decomposition of the forces is shown in SI Figure 4.

We measure the forces for many different values of $\theta_B$ which allows us to integrate the forces in the $\theta_B$ direction over the z coordinate and then map the potential onto $\theta_B$. The total calculation of the bending potential of mean force becomes

$$V(\theta_{B,n}(z_n)) - V(\theta_{B,0}(z_0)) = -\sum_{i=0}^{N-1} \langle F_{\theta_B,i}(z_i)\rangle \left(\langle z_{i+1}\rangle - \langle z_i\rangle\right) \tag{3}$$

### MD Protocol for Calculation of the Bending Potential

The hexamer-hexamer interfaces are solvated in water containing 100 mM NaCl. The system undergoes a short constant pressure, temperature (NPT) equilibration 100 ps with the backbone restrained. The restraints on $Hex_2$ are released while $Hex_1$ is still backbone restrained. Steered MD simulations are then run to create configurations where the proteins are at the many different $\theta_B$ (center of mass z distances) values sampled. This pulling step is done at a rate of 1 Å/ns by a harmonic spring, with a spring constant of 1000 kJ/mol. These N configurations, or ”windows”, are then run in parallel for at least 15 ns to gather the force data necessary for the PMF calculation. For the PduJ we ran 22 windows for 15 and for Supercharged PduA we ran 20 windows for 30ns. Differences in number of windows and run time reflect the complexities of the energy landscapes and an effort to reduce error bars relative to the magnitude of the energies. The error bars are located at the center of each of these ”windows” and thus they also illustrate the window spacing. Error bars are based on sampling error and estimated by splitting the data in different sections (i.e. first third vs. second third vs. last third) and observing the differences in the calculated potential.

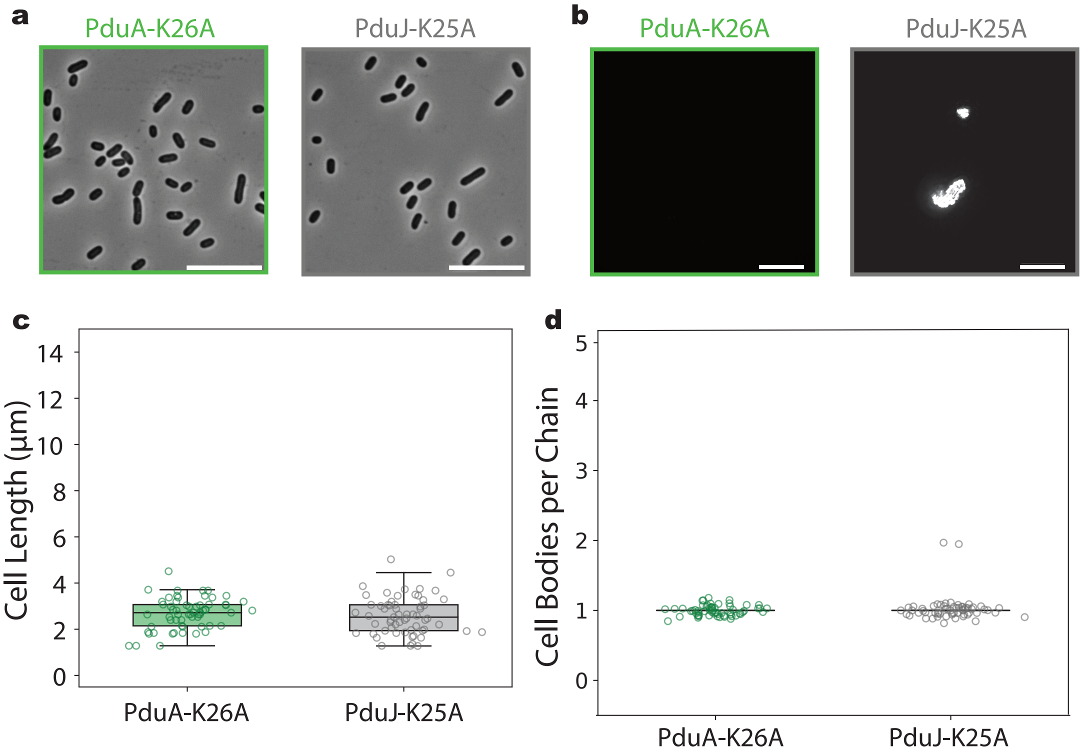


SI Figure 1: **Expression of PduA-K26A and PduJ-K25A assembly-deficient point mutants. a,** Phase contrast image of *E. coli* cell containing assembly-deficient variants of PduA ($K_{26}A$) and PduJ ($K_{25}A$) (scale bar = 10 $\mu$m). Images are captured using phase contrast microscopy. **b,** Cell-free confocal microscopy imaging of assembly-deficient variants of PduA ($K_{26}A$) and PduJ ($K_{25}A$) show no definite structure or just protein aggregation (scale bar = 10 $\mu$m). **c,** Length of cells (defined as end-to-end distance per cell) induced for overexpression ofassembly-deficient variants of PduA ($K_{26}A$) and PduJ ($K_{25}A$). Box plots display distribution of cell lengths including the median, quartiles, mean, and standard deviation across cell population. Measurements were obtained from three biological replicates. **d,** Fraction of linked or elongated cells in total population of cells for assembly-deficient variants of PduA ($K_{26}A$) and PduJ ($K_{25}A$) overexpression in *E. coli* cell. Cells were identified as linked if a clear cleavage furrow was present. An elongated cell, which is $>$ 10 $\mu$m long cells with no cleavage furrow, is counted as a single "linked" cell. Counts were taken from three biological replicates ($>$80 cells counted per strain per replicate).

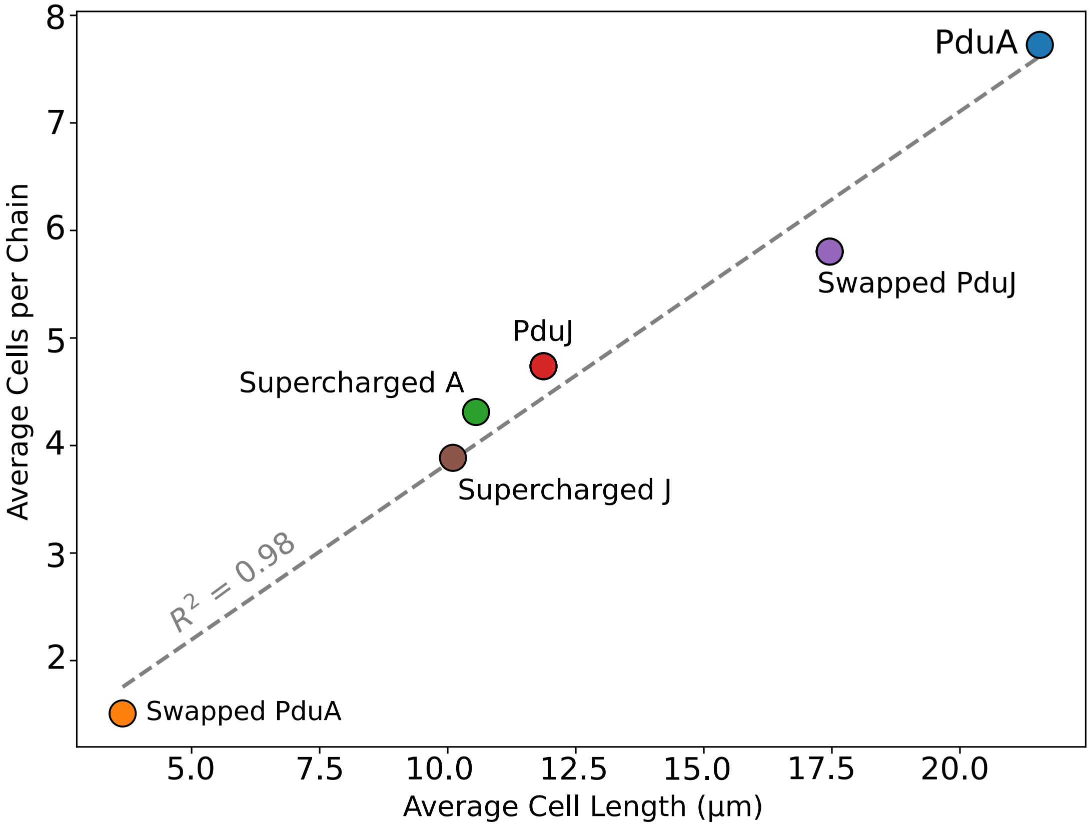


SI Figure 2: **Expression and quantification of Pdu shell protein assembly.** Correlation of average cell length (µm) and fraction of linked or elongated cells shows a positive correlation of $R^2 = 0.98$.

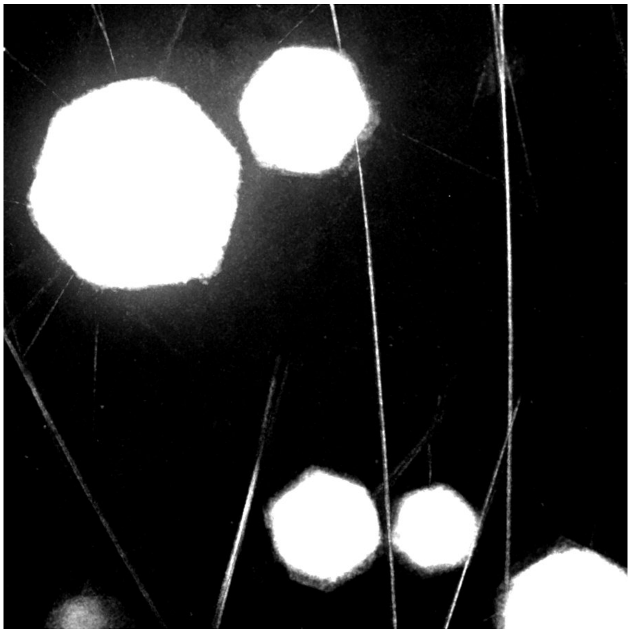

SI Figure 3: **High contrast cell-free image of the WT PduA showcasing both hexagonal sheets and tubular structures.**

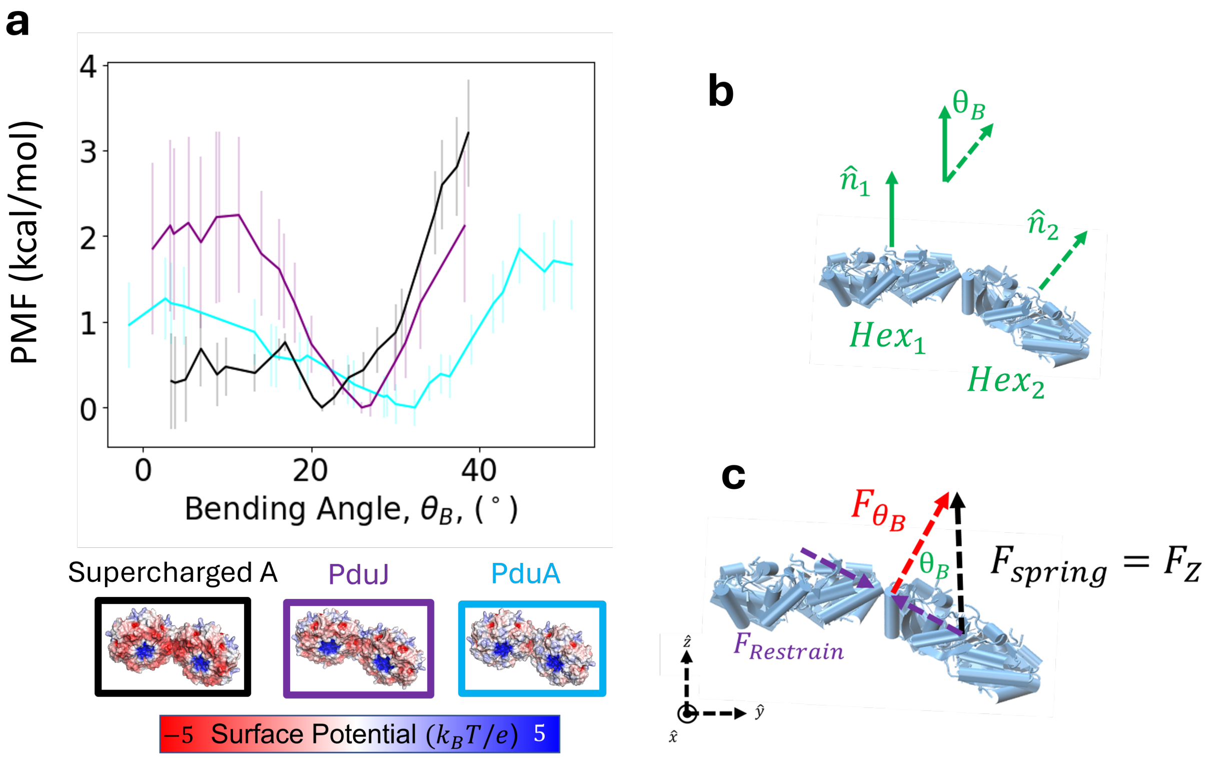


SI Figure 4: **Free-energy cost of bending at Pdu shell protein interfaces a,** Bending potential of mean force calculations for PduA, PduJ, and the Supercharged PduA interfaces. **b,** The definition of the bending angle, $\theta_B$, between Hex$_1$ and Hex$_2$. **c,** The decomposition of the biasing spring force that allows for the calculation of the force in the $\theta_B$ direction.

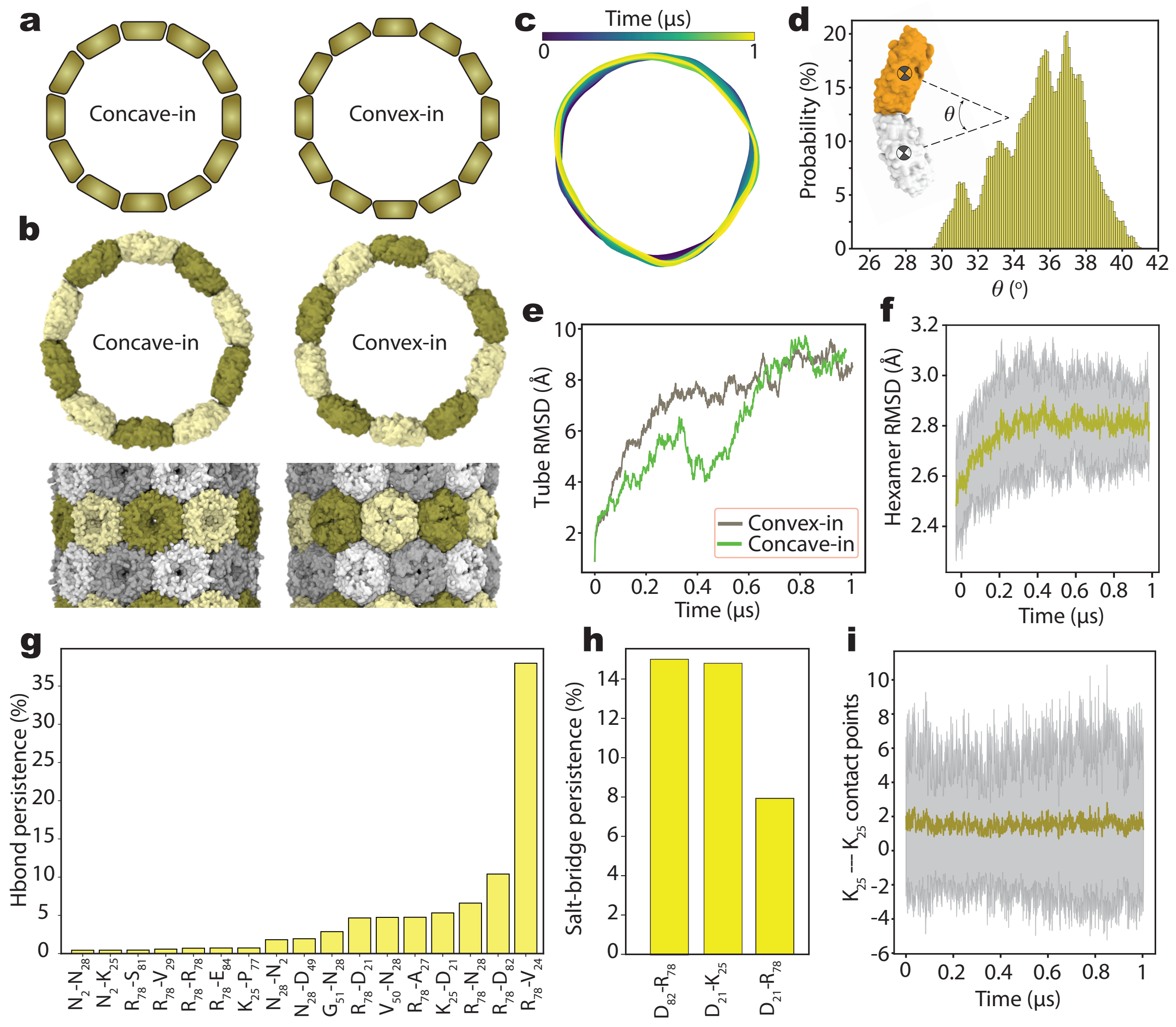


SI Figure 5: **Structural comparison of PduJ tubes: Concave *versus* convex hexamer orientations a,** Schematic representation of a PduJ tube formed from concave-in and convex-in hexamers. PduJ hexamers exhibit a wedge shape when viewed perpendicular to the 6-fold axis. **b,** All-atom model of a 24 nm zigzag nanotube formed from concave-in (right) and convex-in (left) orientations of the hexamers. The top row shows the cross-section of the tube, while the bottom row displays the side view of each tube. **c,** Cross-sectional trajectories of the nanotube from 1 $\mu$s molecular dynamics simulations. The cross-section is visualized by connecting the centers of mass of adjacent hexamers using periodic cubic spline interpolation, resulting in smooth, closed curves. **d,** Distribution of bending angles ($\theta$) between adjacent hexamers for the convex-in configuration of the PduJ tube. **e,** Root Mean Square Deviation (RMSD) of the entire tube over time, calculated using backbone atoms relative to the initial configuration. **f,** RMSD of individual hexamers within the PduJ nanotube, averaged over a 1 $\mu$s trajectory. Error bars represent standard deviations across all hexamers. **g-h,** Persistence of hydrogen bonds (g) and salt bridges (h) between adjacent hexamers, represented as the percentage of simulation time each bond is maintained. Data are averaged over the 1 $\mu$s simulation. **i,** $K_{25}$–$K_{25}$ contact points between PduJ hexamers forming tubular assemblies.

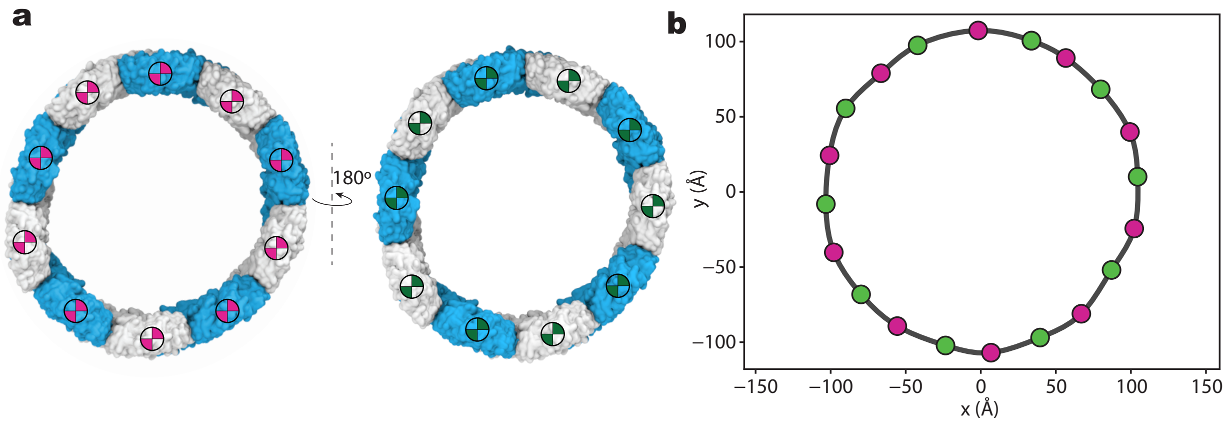


SI Figure 6: **Center-of-mass tracing of the structural pattern in PduJ tube cross section a,** Cross-sectional view of the 24 nm zigzag PduJ tube, with the center-of-mass (CoM) of each hexamer marked. CoMs from alternating rings of the tube are colored magenta and green. **b,** *x*-*y*-plane scatter plot of the hexamer CoMs shown in (a), connected using periodic cubic spline interpolation to form smooth, closed curves.

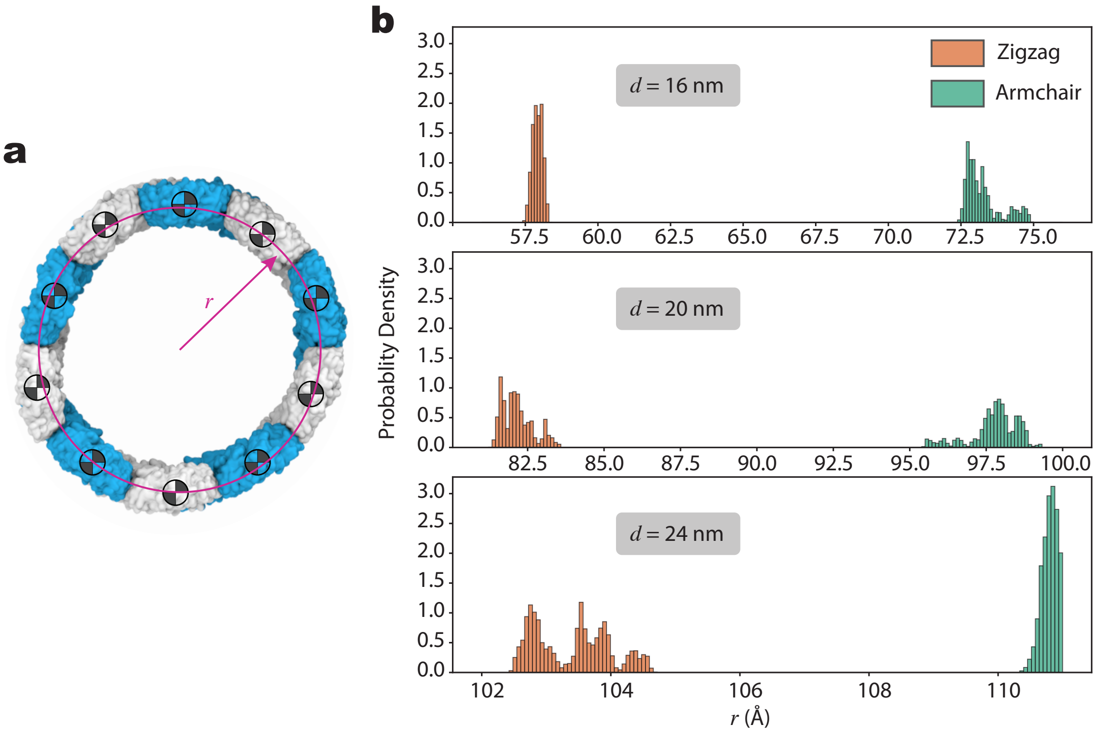


SI Figure 7: **Radius measurements of tubular PduA/PduJ structures.** **a,** The tube radius was determined by fitting a circle to the centers of mass of the hexamers in the *x*–*y* plane (perpendicular to the tube axis). **b,** Distribution of tube radii over the course of the simulations for PduA and PduJ assemblies. Radii were calculated every 10 ps.

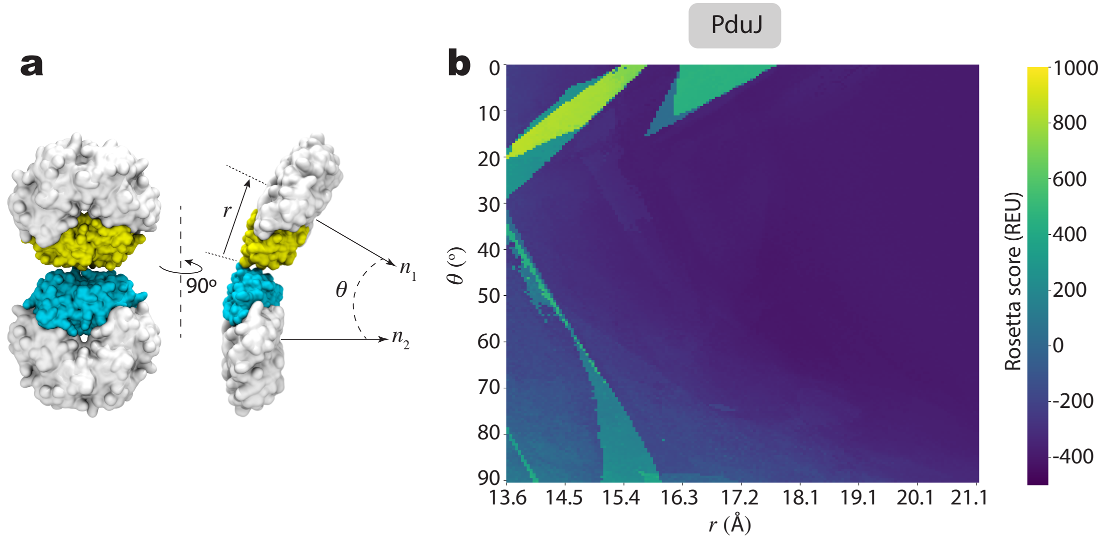


SI Figure 8: **Interaction geometry and energetic landscape of PduJ hexamer assembly a,** Two interacting PduJ hexamers illustrating the bending angle $\theta$ and separation distance between them. The vectors $n_1$ and $n_2$, each normal to the plane of a hexamer and parallel to its 6-fold axis, define the bending angle $\theta$ between the two hexamers. Monomers not included in the simulations are colored white, while simulated monomers are shown in cyan and yellow. **b,** 2D heatmap of Rosetta interface scores as a function of hexamer bending angle (x-axis) and separation distance (y-axis), representing the energetic landscape of hexamer assembly.

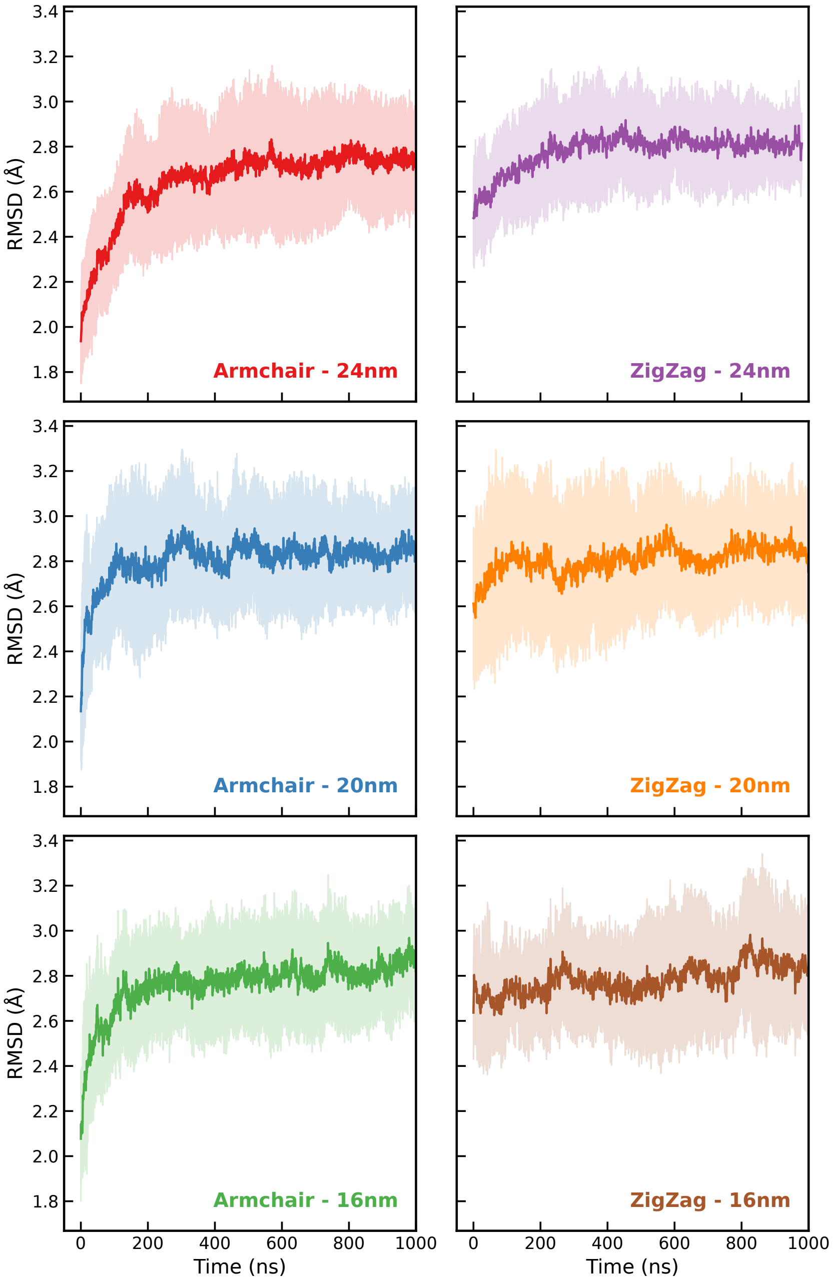


SI Figure 9: **Backbone RMSD of PduJ hexamers forming tubular assemblies.** The plot shows the average backbone RMSD of PduJ hexamers within tubes of varying diameters and chiralities. Shaded regions represent the standard deviation across all hexamers. RMSD values are calculated relative to the crystal structure of PduJ (PDB ID: 5D6V).

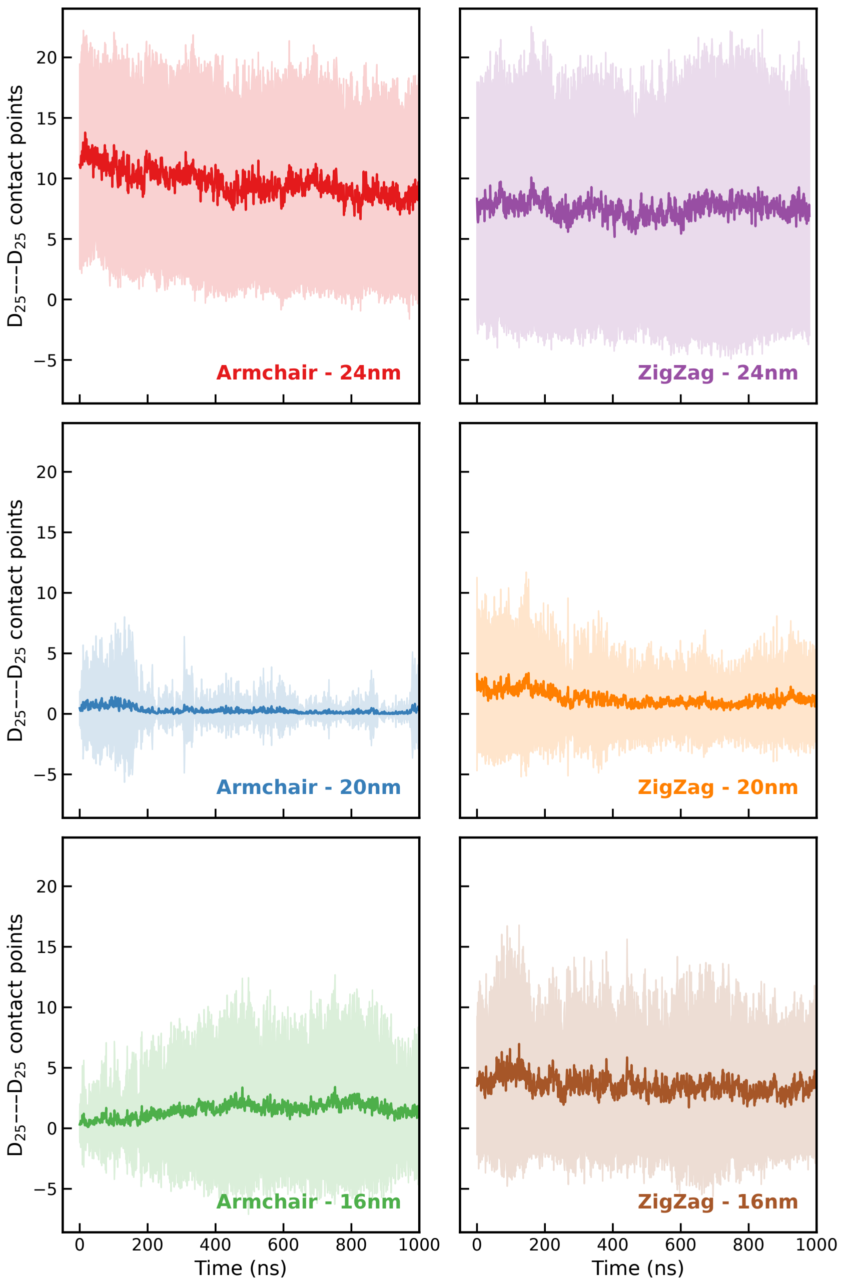


SI Figure 10: **$K_{25}$–$K_{25}$ contact points between PduJ hexamers forming tubular assemblies.** The plot shows the average number of contacts within $K_{25}$–$K_{25}$ paring between adjacent PduJ hexamers in tubes of varying diameters and chiralities. Shaded regions represent the standard deviation across all hexamer pairs. Contact points are defined as atom pairs within 3.5 Å of each other within the $K_{25}$ residues.

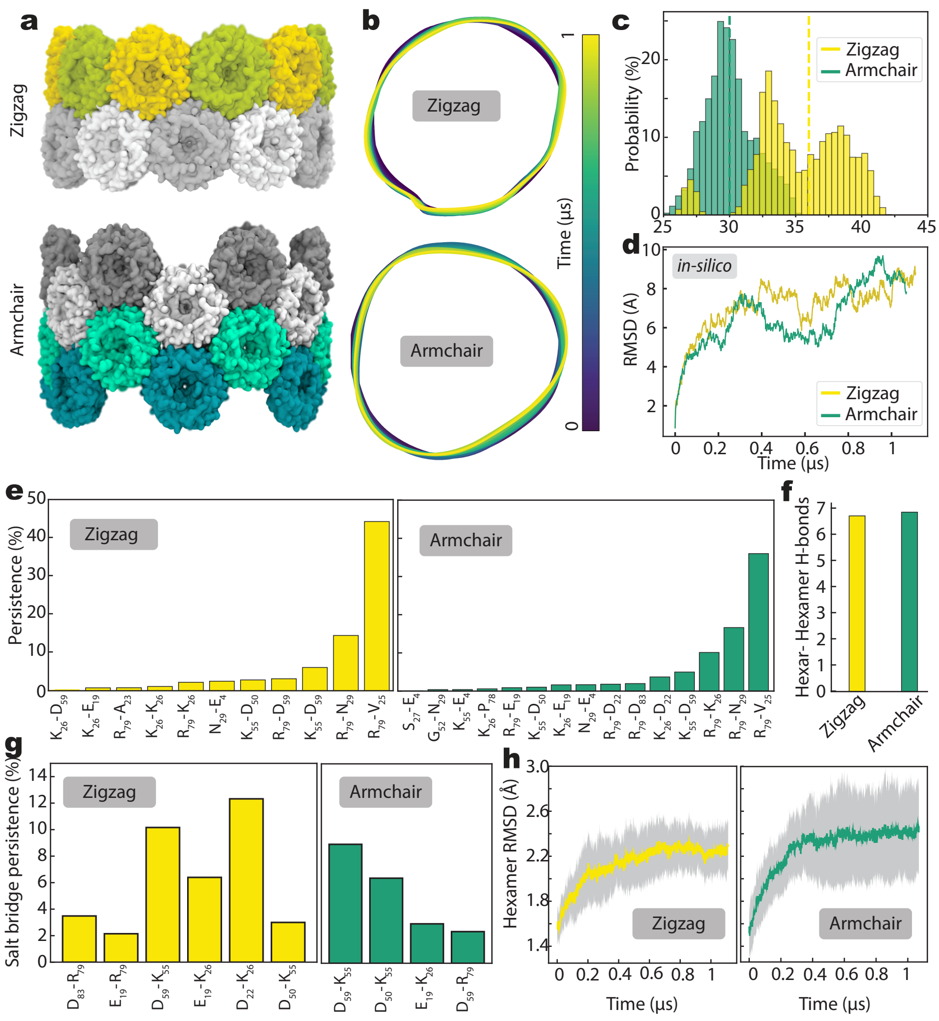


SI Figure 11: ***In silico* model of a 24 nm PduA nanotube. a,** All-atom representation of a 24nm PduA nanotube, exhibiting both armchair (green) and zigzag (yellow) chiralities. The tube is solvated in a hexagonal water box (not shown), which defines the periodic simulation cell. **b,** Cross-sectional trajectories of the nanotube over 1$\mu$s molecular dynamics simulations for both chiral forms. The cross-section is visualized by connecting the centers of mass of adjacent hexamers using periodic cubic spline interpolation, resulting in smooth, closed curves. **c,** Distribution of bending angles ($\theta$) between adjacent hexamers for the armchair (green) and zigzag (yellow) configurations. The angle $\theta$ is defined between vectors from each hexamer's center of mass to the tube center (top-right inset). The dashed line marks the ideal bending angle for a perfect circle, derived from the tube radius and hexamer count. **d,** Root Mean Square Deviation (RMSD) of the entire tube over time, calculated using backbone atoms relative to the initial configuration. **e–f,** Persistence and count of hydrogen bonds between adjacent hexamers, represented as the percentage of simulation time each bond is maintained. Data are averaged over the 1 $\mu$s simulation. **g,** Average salt bridge persistence per monomer across the simulation. **h,** RMSD of individual hexamers within the PduJ nanotube, averaged over a 1 $\mu$s trajectory. Error bars denote standard deviations across all hexamers. RMSD values are referenced to the crystal structure of the PduJ hexamer (PDB ID: 3NGK).

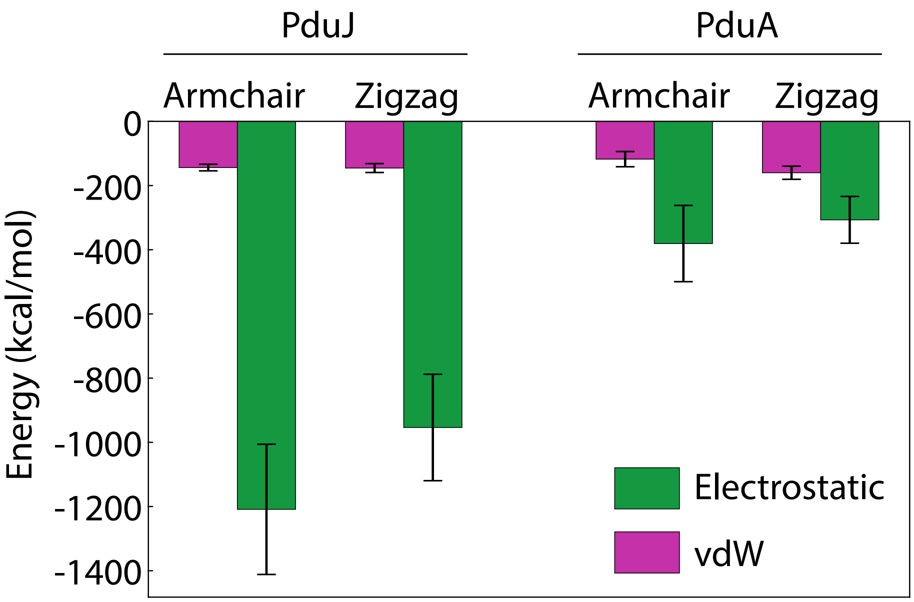


SI Figure 12: **Non-bonded interaction energies of a hexamer within the tubular assembly.** Electrostatic and van der Waals (vdW) interaction energies between an individual hexamer and the remainder of the tube in the assembled configuration. Energies are averaged over all symmetry-equivalent hexamers and over the final 200 ns of the production simulations. Error bars represent the standard error of the mean.

#### SI Table 1: Summary of PduA and PduJ variant testing across expression environments.

| Variant name | Heterologous Overexpression | Cell-Free Protein Synthesis | In Vivo |
|---|---|---|---|
| **PduA variants** | | | |
| WT PduA | pBAD33t-*pduA*-FLAG | pJL1-PduA-FLAG | $\Delta pduJ$ |
| Swapped PduA | pBAD33t-*pduA*-E4N-R66S-K86A-FLAG | pJL1-PduA-E4N-R66S-K86A-FLAG | $\Delta pduA :: pduA$-E4N-R66S-K86A |
| Supercharged PduA | pBAD33t-*pduA*-E4K-R66E-K86E-FLAG | pJL1-PduA-E4K-R66E-K86E-FLAG | $\Delta pduA :: pduA$-E4K-R66E-K86E $\Delta pduJ$ |
| **PduJ variants** | | | |
| WT PduJ | pBAD33t-*pduJ*-FLAG | pJL1-PduJ-FLAG | $\Delta pduA :: pduJ$ $\Delta pduJ$ |
| Swapped PduJ (N3E, S65R, A85K) | pBAD33t-*pduJ*-N3E-S65R-A85K-FLAG | pJL1-PduJ-N3E-S65R-A85K-FLAG | $\Delta pduA :: pduJ$-N3E-S65R-A85K $\Delta pduJ$ |
| Supercharged PduJ (N3K, S65E, A85E) | pBAD33t-*pduJ*-N3K-S65E-A85E-FLAG | pJL1-PduJ-N3K-S65E-A85E-FLAG | $\Delta pduA :: pduJ$-N3K-S65E-A85E $\Delta pduJ$ |

SI Table 2: Structure parameters and atom counts of the simulated PduJ systems.

| | **Chirality** (n,m) | **d** (nm) | **Hexamers** (per Ring) | $\theta$ ($^o$ deg) | **Finite system** (atoms) | **infinite system** (atoms) |
|---|---|---|---|---|---|---|
| **Armchair** | (6,6) | 24 | 12 | 30 | 2,758,070 | 110,6222 |
| | (5,5) | 20 | 10 | 36 | 2,467,484 | 957,969 |
| | (4,4) | 16 | 8 | 45 | 1744091 | 675,347 |
| **Zigzag** | (10,0) | 24 | 10 | 36 | 2,128,561 | 939,960 |
| | (8,0) | 8 | 10 | 45 | 1,632,112 | 717,708 |
| | (6,0) | 60 | 6 | 60 | 1,091,466 | 480,396 |

SI Table 3: Plasmids used in this study.

| Name | Plasmid | Origin | Resistance |
|---|---|---|---|
| pSIM6 | $\lambda$ Red system repressed by cI857 | pSC101 repA$^{ts}$ | Ampicillin |
| CMJ069 | pBAD33t-ssD-GFPmut2 | p15A | Chloramphenicol |
| pCEM028 | pBAD33t-ssD-GFPmut2 | p15A | Chloramphenicol |
| pCEM419 | pBAD33t-PduA-FLAG | p15A | Chloramphenicol |
| pCEM420 | pBAD33t-PduJ-FLAG | p15A | Chloramphenicol |
| pCEM112 | pBAD33t-PduA-E4K-R66E-K86E-FLAG | p15A | Chloramphenicol |
| pCEM113 | pBAD33t-PduA-E4N-R66S-K86A-FLAG | p15A | Chloramphenicol |
| pCEM114 | pBAD33t-PduJ-N3K-S65E-A85E-FLAG | p15A | Chloramphenicol |
| pCEM115 | pBAD33t-PduJ-N3E-S65R-A85K-FLAG | p15A | Chloramphenicol |

SI Table 4: Primers used in this study. Homology region is underlined.

| Name | Purpose | Description | Sequence* |
|---|---|---|---|
| NWKo418 | Recombineering | Amplify *pduA* with homology at *pduA* For | GCATCTTCTTATAGTCCCAACTATC GGAACACTCCATGCGAGGTCTTT ATGCAACAAGAAGCACTAGGAATG |
| NWKo419 | Recombineering | Amplify *pduA* with homology at *pduA* Rev | GGGCAATCACCTGCGCCATGATCTG TTCCACCAGCTCATTGCTGCTCATT GGCTAATTCCCTTCGGTAAGA |
| oASG001 | Recombineering | Amplify *PduA*-E4K-R66E-K86E-FLAG with homology at *pduA* For | GCATCTTCTTATAGTCCCAACTATC GGAACACTCCATGCGAGGTCTTT ATGCAACAAAAAGCACTAGGAATG |
| oASG002 | Recombineering | Amplify *PduA*-E4N-R66S-K86A-FLAG with homology at *pduA* For | GCATCTTCTTATAGTCCCAACTATC GGAACACTCCATGCGAGGTCTTT ATGCAACAAAACGCACTAGGAATG |
| oASG003 | Recombineering | Amplify *PduJ*-N3K-S65E-A85E-FLAG with homology at *pduA* For | GCATCTTCTTATAGTCCCAACTATC GGAACACTCCATGCGAGGTCTTT ATGAATAAGGCACTGGGACTGG |
| oASG004 | Recombineering | Amplify *PduJ*-N3E-S65R-A85K-FLAG with homology at *pduA* For | GCATCTTCTTATAGTCCCAACTATC GGAACACTCCATGCGAGGTCTTT ATGAATGAAGCACTGGGACTGG |
| oASG005 | Recombineering | Amplify *PduJ*-N3K-S65E-A85E-FLAG with homology at *pduA* Rev | GGGCAATCACCTGCGCCATGATCTG TTCCACCAGCTCATTGCTGCTCATT GGCTAATTCCCTTCGGTAAGATTTT TTCTACATCGGTGTGAGGGCGTTAG GCTGATTTCGGTAAAATTTTC |
| oASG006 | Recombineering | Amplify *PduJ*-N3E-S65R-A85K-FLAG with homology at *pduA* Rev | GGGCAATCACCTGCGCCATGATCTG TTCCACCAGCTCATTGCTGCTCATT GGCTAATTCCCTTCGGTAAGATTTT TTCTACATCGGTGTGAGGGCGTTAG GCTGATTTCGGTAAAATTTCC |
| oCEM252 | Golden gate amplification | PduA-E4K-R66E-K86E mutagenesis For | ATTAGGTCTCACATGCAACAAAAAGCA |
| oCEM253 | Golden gate amplification | PduA-FLAG mutagenesis Rev | TAATGGTCTCATTTACTTGTCATCGTCA |
| oCEM254 | Golden gate amplification | PduA-E4N-R66S-K86A mutagenesis For | ATTAGGTCTCACATGCAACAAAACGCA |
| oCEM255 | Golden gate amplification | PduJ-N3K-S65E-A85E-mutagenesis For | ATTAGGTCTCACATGAATAAGGCACTG |
| oCEM256 | Golden gate amplification | PduJ-N3E-S65R-A85K mutagenesis For | ATTAGGTCTCACATGAATGAAGGCACTG |

*Homology region is underlined.

(17) Mills, C. E.; Waltmann, C.; Archer, A. G.; Kennedy, N. W.; Abrahamson, C. H.; Jackson, A. D.; Roth, E. W.; Shirman, S.; Jewett, M. C.; Mangan, N. M., et al. Ver-

tex Protein PduN Tunes Encapsulated Pathway Performance by Dictating Bacterial Metabolosome Morphology. *Nature Communications* **2022**, *13*, 3746.

(18) Waltmann, C.; Shrestha, A.; Olvera de la Cruz, M. Patterning of multicomponent elastic shells by gaussian curvature. *Physical Review E* **2024**, *109*, 054409.

(19) Schrödinger, LLC, The PyMOL Molecular Graphics System, Version 1.8.
